# Evolutionary hotspots of structural variation drive inter-individual differences in the expression of fusion transcripts in the human brain

**DOI:** 10.1101/2025.03.11.642572

**Authors:** Colette Moses, Angelica Vanini, Maria Chiara Bocchi, Luca Wagner, Isis van Loenen, Brooke L. Latour, Nael Nadif Kasri, Frank M. J. Jacobs

## Abstract

While most protein-coding regions in our genome are highly conserved, phenotypic variation largely arises from differences in gene transcription, splicing, and translation. Intra-gene splicing variations are a known source of inter-individual differences. However, inter-gene splicing events, which generate fusion transcripts—RNA molecules combining exons from multiple distinct genes—are less explored. Fusion transcripts are well-studied in cancer, but their expression and inter-individual variability in normal human tissues has not been thoroughly investigated. We conducted a genome-wide fusion transcript analysis in postmortem human brain tissues from 276 individuals, identifying 717 distinct fusion transcripts. Many had protein coding potential, further supported in some cases by ribosome profiling data. Fusion transcripts that were present in only part of the human population were predominantly located within segmental duplication (SD) “hotspots”; highly copy number-variable genomic regions linked to neurodevelopmental and neurodegenerative diseases. None of these variable fusion transcripts were present in chimpanzee or rhesus macaque, suggesting they are human-specific. We confirmed that SD-derived fusion transcripts arise from structural genomic rearrangements in the human population, previously associated with neurodevelopmental and neurodegenerative disease risk. Inter-individual variation in fusion transcript expression represents an overlooked source of genetic diversity, with potential to contribute to differences in disease susceptibility.

## Introduction

Comparative genomics has revealed multiple ways in which genome evolution gives rise to species-specific gene expression profiles^1^. Genomic changes can impact spatial and temporal dynamics of gene expression, while alternative splicing can produce novel transcript variants and protein isoforms, even in genes well-conserved at the exon level^2–5^. Additionally, entire genes can be lost or gained throughout evolution, potentially disrupting or innovating important gene expression networks^6^. A special form of alternative splicing can generate fusion transcripts, where exons from two or more distinct genes are combined^7,8^. These are often the result of “readthrough transcription” of adjacent or nearby genes on the same strand, a process whereby both genes are transcribed into a single pre-mRNA (**Figure 1A**). Splicing then leads to a mature transcript composed of exons originating from multiple genes. While readthrough transcription usually occurs at low levels, some fusion transcripts arising from this phenomenon are highly expressed and generate functional fusion proteins^7,9,10^.

**Figure 1.**
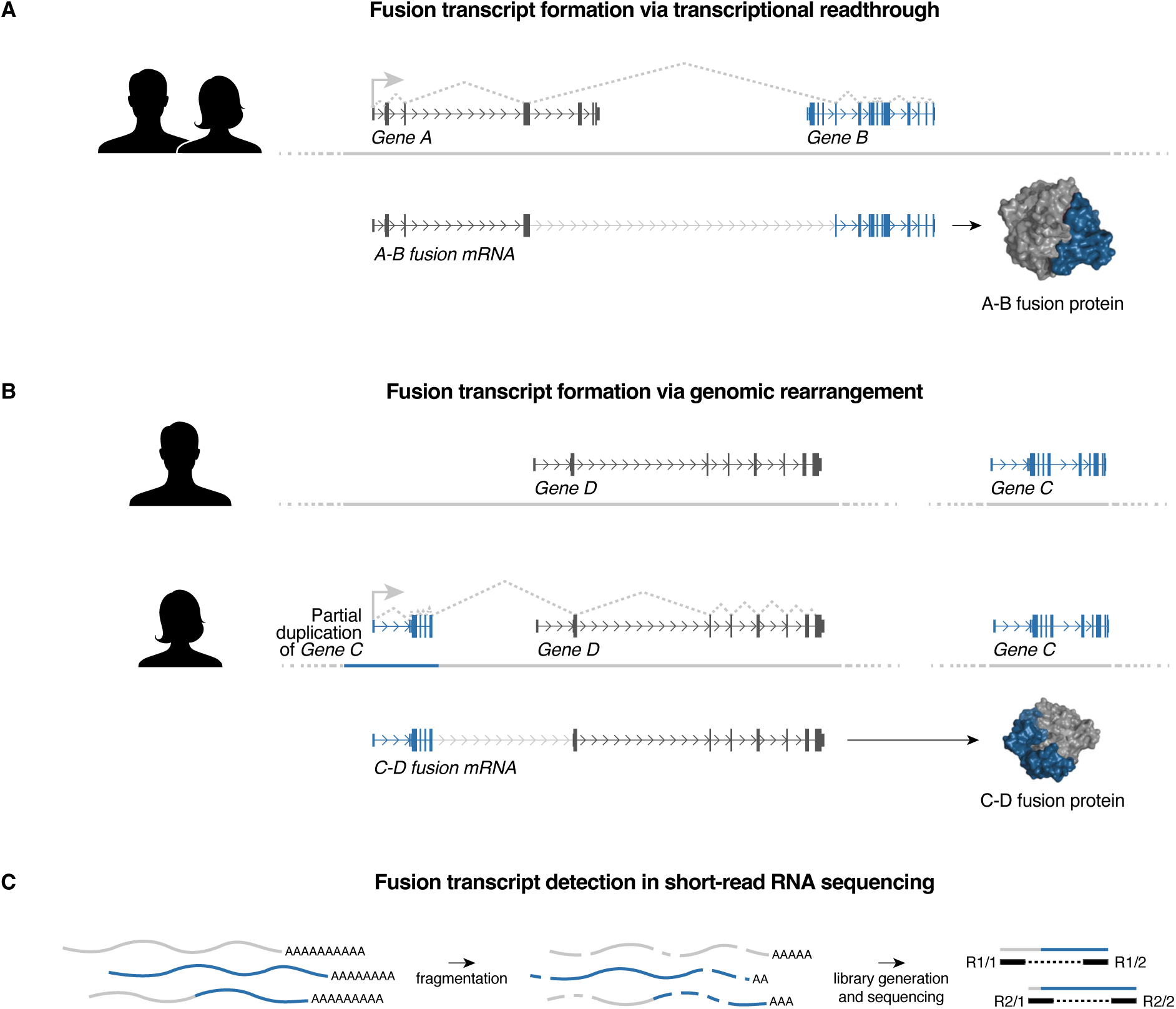
Fusion transcript formation and detection. **A**: Fusion transcripts can form between adjacent genes on the same strand by the process of transcriptional readthrough. Transcription starts in the upstream gene and instead of terminating at the end of the gene, continues through the intergenic region and into the downstream gene. The pre-mRNA is then spliced, resulting in a fusion mRNA containing exons from both original genes. **B**: Fusion transcripts can also arise as a result of genomic rearrangements, such as insertions, deletions, inversions and translocations. In one individual, two genes may be located in disparate parts of the genome. However, a genomic rearrangement in another individual can place genetic elements in proximity to one another. This then allows a fusion transcript to form through the transcriptional readthrough mechanism. **C**: Fusion transcripts are detected in short-read RNA sequencing data by identifying reads or read pairs that map simultaneously to more than one annotated gene.

More sporadically, fusion transcripts can arise as a result of genomic rearrangements such as inversions, insertions, deletions and duplications^11^. These rearrangements can bring into proximity previously disparate genes or exons, causing them to be transcribed together into a single RNA molecule (**Figure 1B**). Certain human genome regions are inherently genetically unstable due to the accumulation of segmental duplications (SDs) in the last 18-14 million years, resulting in a highly complex and repetitive genomic structure^12–14^. Ongoing copy number variations (CNVs) at these locations often cause neurological disorders such as autism spectrum disorder, developmental delay, intellectual disability and other neurodevelopmental conditions^15–18^. Regions rich in segmental duplications may be prone to generating fusion transcripts by the accidental combination of two or more functionally unrelated partial gene copies in the locus. In addition, partial segmental duplication of genes can lead to the emergence of truncated transcripts and proteins, which can adopt novel functions, often by competing or interacting with the parental protein^18–20^.

Fusion transcripts, whether they arise from transcriptional readthrough or novel chromosomal rearrangements, increase the diversity of the transcriptome. Some possess different expression patterns than their parental genes and give rise to novel protein isoforms with specific functions^7,9,21–23^. For instance, a fusion transcript formed from the *CTNNBIP1* and *CLSTN1* genes was recently implicated in the proliferation of neuronal progenitors during human corticogenesis^24^. This transcript is specifically enriched in outer radial glia, absent in mice, and its downregulation leads to precocious neuronal differentiation, suggesting a role in human brain development and evolution. Other examples of fusion genes with known functions include the chemokine receptor *CCL14-CCL15*, the luteinizing hormone receptor *SBLF-ALF*, and the drug-metabolising enzyme *CYP2C18-CYP2C19*^25–27^.

Most genes in our genome are believed to have undergone some level of inter-gene recombination in our evolutionary past. Modern multi-domain proteins likely evolved from simpler, single-domain proteins through genomic and transcriptomic fusion events^28^. We hypothesise that this evolutionary phenomenon continues today, with fusion transcripts generated through transcriptional readthrough or underlying genomic rearrangements serving as a source of evolutionary innovation in our genome, transcriptome and proteome. Unsurprisingly, fusion transcripts are commonly observed in situations of pathological genomic instability such as cancer^29,30^. However, the existence and expression of fusion transcripts in healthy tissues, and inter-individual variation of their expression patterns in the human population, remain largely elusive. In this study, we set out to identify and characterise fusion transcript expression across hundreds of individuals in post mortem brain tissues to create a genome-wide catalogue of transcriptional fusion events and assess inter-individual variation in fusion transcript expression. Our work reveals that fusion transcripts can arise in highly duplicated loci as a direct result of evolutionary genomic rearrangements, so recent that they are present in some but not all individuals in the human population. Given that these loci are inherently prone to both pathological and benign copy number variations, the presence or absence of fusion transcripts may vary widely, with unexplored functional consequences for physiological and pathophysiological processes in each individual.

## Results

### Fusion transcripts are widely expressed in the human brain and highly variable between individuals

We set out to analyse fusion transcript expression in human brain tissue, as many genes that play critical roles in the nervous system have previously been shown to reside within genomic rearrangement hotspots^31^. To identify fusion transcripts genome-wide, we analysed and compared transcriptome data from postmortem temporal cortex tissue of 276 individuals. We used the program FusionCatcher^32^ to search for transcripts based on short-read RNA sequencing (RNA-seq) reads and read pairs that map simultaneously to more than a single annotated gene (**Figure 1C**). Fusion transcripts that were previously identified in healthy tissue databases were not discarded. After filtering, 717 fusion transcripts remained, of which 94% were intrachromosomal (**Figure 2A**; **Supplemental Figure 1A**). After identifying this set of fusion transcripts with FusionCatcher, we used bowtie2 to map reads directly to the fusion junction sequences. This targeted approach was able to capture a greater number of supporting reads than standard FusionCatcher analysis alone.

**Figure 2.**
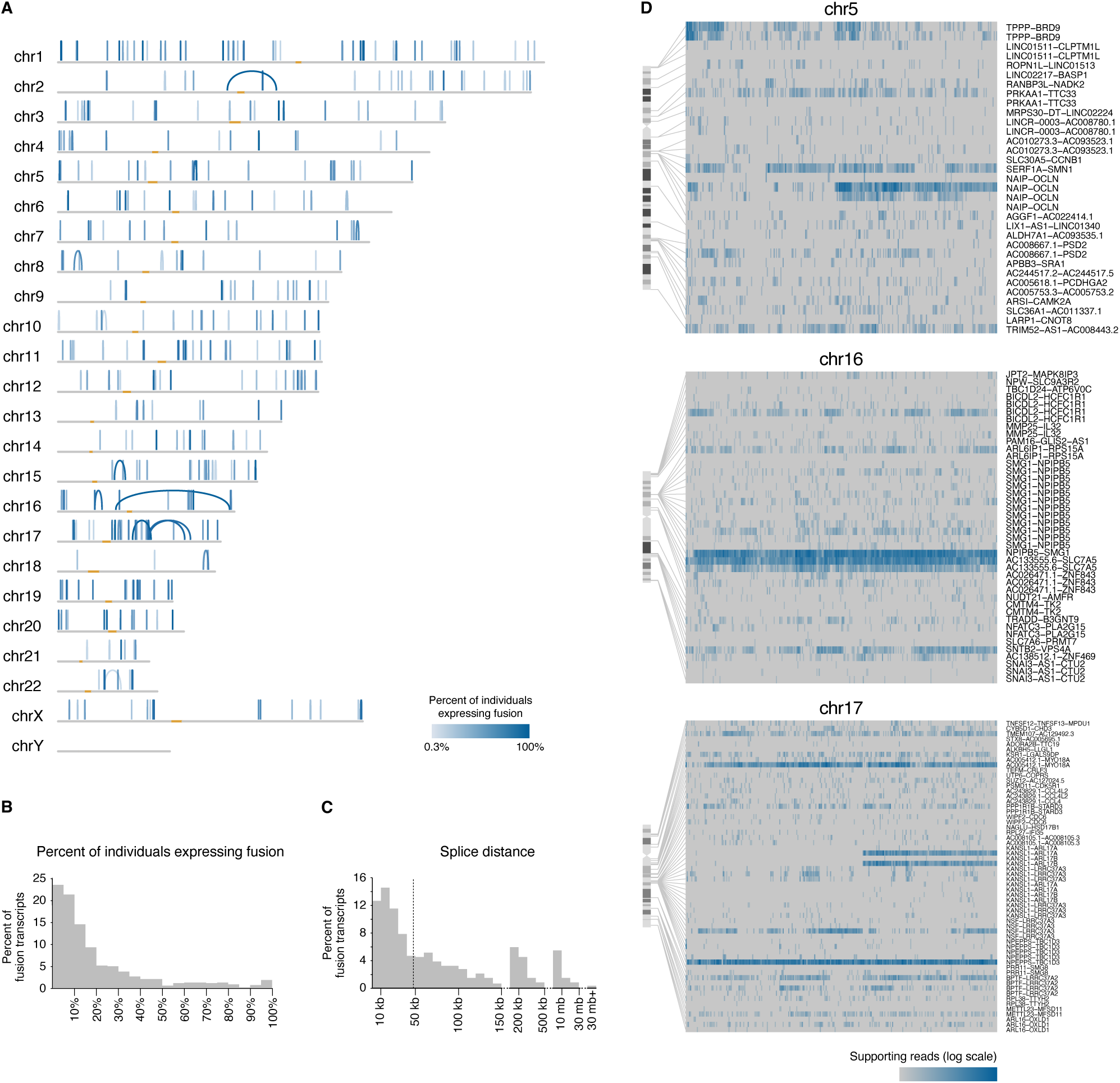
Genome-wide fusion transcript analysis in human brain. **A**: Representation of 717 fusion transcripts detected in two or more individuals in postmortem human brain RNA-seq data from 276 individuals. Each blue arc connects the upstream and downstream fusion points of a single detected fusion transcript. For fusion transcripts where the upstream and downstream transcripts are a short distance apart, the arc appears as a vertical line. The depth of colour of each arc corresponds to the percentage of individuals in which the fusion transcript was detected. Centromeres are depicted in orange. Only intrachromosomal fusions are shown; interchromosomal fusions are displayed in Supplemental Figure 1. **B**: Percent of fusion transcripts detected in differing numbers of individuals. **C**: Percent of fusion transcripts with different splice distances. Splice distance refers to the genomic distance between upstream and downstream fusion points. Median splice distance is shown with a dotted line. **D**: Heatmaps displaying fusion transcripts detected per individual on each chromosome (4 representative chromosomes are shown; remaining chromosomes are shown in **Supplemental Figure 2**). Rows represent different fusion transcripts, and columns represent individuals. Depth of colour is proportional to the number of RNA-seq reads supporting the fusion.

There was a high degree of variation in the number of individuals expressing each fusion transcript. Some transcripts were expressed in almost all individuals, while most were detected in less than 20% of individuals (**Figure 2B**). On average, each individual showed expression of 20% (142/717) of fusion transcripts (**Supplemental Figure 1**). The median splicing distance between the fusion points of the upstream and downstream parental genes was 47.7 kilobases (**Figure 2C**). Seventy percent (502/717) of fusions occurred between immediately adjacent genes, and of these, 98% (492/502) occurred in the direction that would be predicted from standard readthrough transcription of nearby genes. However, there was a substantial number of candidate fusions occurring between very distant genes or contrary to the expected direction of transcriptional readthrough, suggesting that these transcripts were arising from genomic rearrangements, trans-splicing events, or from sequences that were not annotated correctly in the human reference genome.

We further validated the 717 fusion transcripts in long-read RNA-seq data from cerebellum and cortex tissue of six individuals^33,34^. While long-read sequencing data are not available for as many individuals, this technique provides more detailed insight into the full-length structures and genomic locations of fusion transcripts. Seventy-four percent (527/717) of fusion transcripts we previously identified in short-read RNA-seq were validated by corresponding long reads mapping to the fusion transcript junction in one or more individuals. This assessment supports that the majority of the 717 identified fusion transcripts are genuine fusions, rather than false positives arising from mapping errors or other artefacts from short-read data. Long-read sequencing revealed that on average, each individual expressed 39% (281/717) of fusion transcripts that had originally been identified in the short-read dataset (**Supplemental Figure 3A**). It should be noted that because only six long-read datasets were used to verify fusion transcripts detected in 276 individuals, the actual proportion of the 717 fusion transcripts that could be validated by long read sequencing may be higher if more transcriptomes are analysed in the future.

### Fusion transcripts potentially encode novel protein isoforms

We next determined what proportion of fusion transcripts could potentially code for fusion proteins. The ability of a fusion transcript to produce a functional fusion protein depends on several factors, including whether the fusion involves protein-coding or non-coding genes and the location of the fusion junction between the upstream and downstream genes. Examples of different fusion possibilities are displayed in **Figure 3A-D**. Fusion transcripts occurring between the coding sequence (CDS) of two protein-coding genes can be fused in-frame (**Figure 3A**) or out-of-frame, potentially encoding a full-length fusion protein or a truncated protein, respectively. Additionally, some fusions that do not arise from two CDS may nevertheless contain open reading frames. For example, the CDS of an upstream gene may fuse to the 5’ untranslated region (UTR) of a downstream gene and continue into the CDS of the downstream gene in-frame (**Figure 3B**). Fusions between the 5’ UTR of an upstream gene and the CDS of a downstream gene could potentially utilise an alternative start site for translation (**Figure 3C**). Additionally, fusions of long non-coding RNAs (lncRNAs) also have the potential to produce novel fusion transcripts, in combination with protein-coding genes (**Figure 3D**) or other lncRNAs.

**Figure 3.**
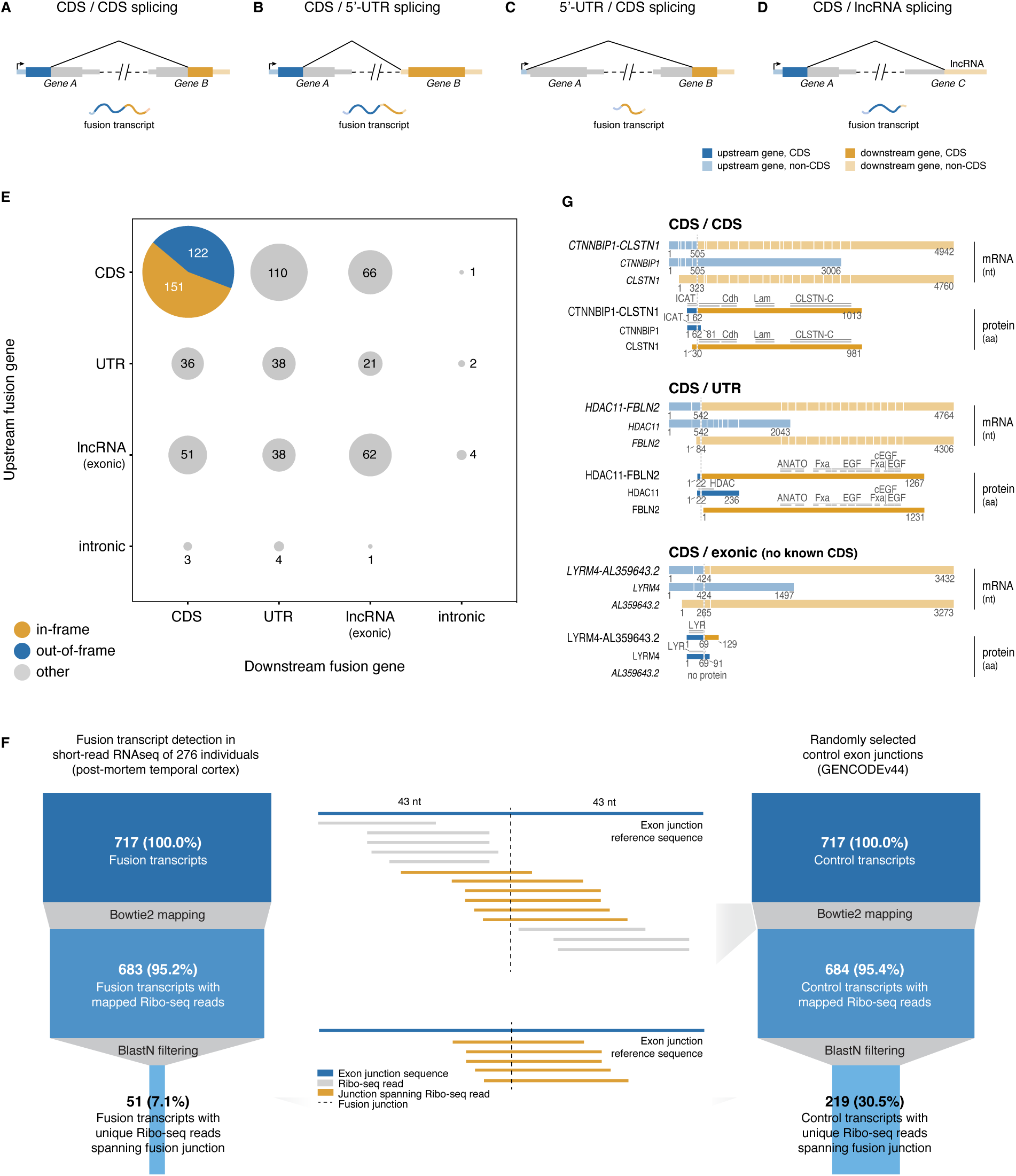
Fusion transcripts encode potentially functional proteins and are detected in ribosome profiling data. **A-D**: Examples of common types of fusion transcripts formed from fusions at various locations within the upstream and downstream genes. CDS: coding sequence; UTR: untranslated region; lncRNA: long non-coding RNA. **E**: Number of fusion transcripts with different protein-coding potential detected in the postmortem human brain RNA-seq dataset, displayed according to combination of upstream and downstream fusion locations (n=717; 7 fusion transcripts had unknown protein-coding potential and are not displayed). UTR refers to the 5’ or 3’ untranslated region of a protein-coding gene. **F**: Numbers of fusion transcripts and randomly selected control transcripts detected in ribosome profiling (Ribo-seq) data. Results from one individual genetic background are shown here; data from 5 more individual genetic backgrounds are provided in Supplemental File 1. **G**: Examples of fusion transcripts and encoded proteins falling within several categories defined in panel A, which were also detected in ribosome profiling data. Location of the fusion junction is indicated with a dotted line and functional domains are labelled.

We characterised the set of 717 fusion transcripts based on the nature of the upstream and downstream fusion genes and found that 38% (273/717) of fusion events resulted in CDS/CDS merging (**Figure 3E**). Of the CDS/CDS fusions, 55% (151/273) fused in-frame. These transcripts have the potential to be translated into a fusion protein that is a chimera of the two parental proteins. The remaining 45% (122/273) produced out-of-frame transcripts, likely encoding truncated proteins. Other highly represented categories were fusions between the CDS of the upstream gene and the UTR of a downstream protein-coding gene (15% of total, 110/717), or an exon of a downstream non-coding gene (9% of total, 66/717). These categories could lead to truncated or novel proteins, depending on the specific case.

To determine whether any fusion transcripts showed evidence of active protein translation, we analysed ribosome profiling (Ribo-seq) data^35^. It should be noted that even for normal genes, detection in Ribo-seq is limited by low coverage and short read length. Detection of fusion transcripts is even more challenging because Ribo-seq reads can only unambiguously support fusion transcripts if they span the specific exon-exon fusion junction, rather than any location along the transcript. Despite these technical limitations, 7% of fusion transcripts (51/717) showed expression in ribosome footprints from one individual genetic background, compared to 31% (219/717) of randomly selected control genes from GENCODEv44 (**Figure 3F**; analysis for five more individuals is shown in **Supplemental File 1**). While 7% of fusion transcripts may seem like a small fraction, it is relevant that any given individual expresses only a subset of all 717 fusion transcripts (39% based on long read sequencing data). In addition, for the control set of annotated genes, expected to be more stably expressed across the population than fusion transcripts, only 31% of exon-exon junctions were identified in Ribo-seq, highlighting the inherent limitations of ribosome-transcript interactions and the ribosome profiling technique. A closer examination of specific fusion transcripts detected in Ribo-seq revealed their potential to generate novel chimeric or truncated proteins (**Figure 3G**). Overall, these findings suggest that some fusion transcripts are actively translated, and many others may go undetected due to technical challenges in identifying fusion junction sequences in Ribo-seq data.

### Some fusion transcripts exhibit pronounced inter-individual variation, while others are readthrough events common to most or all individuals

We next assessed whether the fusion transcripts identified through our bioinformatic pipeline could be validated in independent RNA samples. We selected induced pluripotent stem cell lines from 9 individuals in the Hipsci database^36^, for which both high coverage RNA-seq and whole genome sequencing (WGS) data are available. We performed RT-PCR for 20 selected fusion transcripts and cross-referenced these results with the expected expression from FusionCatcher analysis performed on RNA-seq data from the same cell lines. We successfully validated 18/20 of these fusions by RT-PCR (**Figure 4**; **Supplemental Figure 4**). Among the validated fusion transcripts were those arising from adjacent genes, as well as from genes located far apart in the reference genome. Four fusion transcripts showed differential expression between individuals, all of which arose from distant genes in the reference genome (**Figure 4A**). These transcripts (*NAIP-OCLN*, *KANSL1-ARL17A/B*, *KANSL1-LRRC37A3* and *NSF-LRRC37A3*) were located within known alternate haplotype regions of the human genome, suggesting they arose from diverse genomic structural arrangements in the nine individuals profiled. In contrast, fusion transcripts that arose from readthrough transcription of adjacent genes were expressed in all individuals based on RT-PCR, even when they had not been detected in all individuals by RNA-seq (**Figure 4B**; **Supplemental Figure 4**). This implies that the sensitivity of FusionCatcher detection of fusion transcripts in RNA-seq data is suboptimal, and in particular the set of transcriptional readthrough-derived fusion transcripts may be expressed more widely in the human population than what is predicted based on the bioinformatic approaches used in this study.

**Figure 4.**
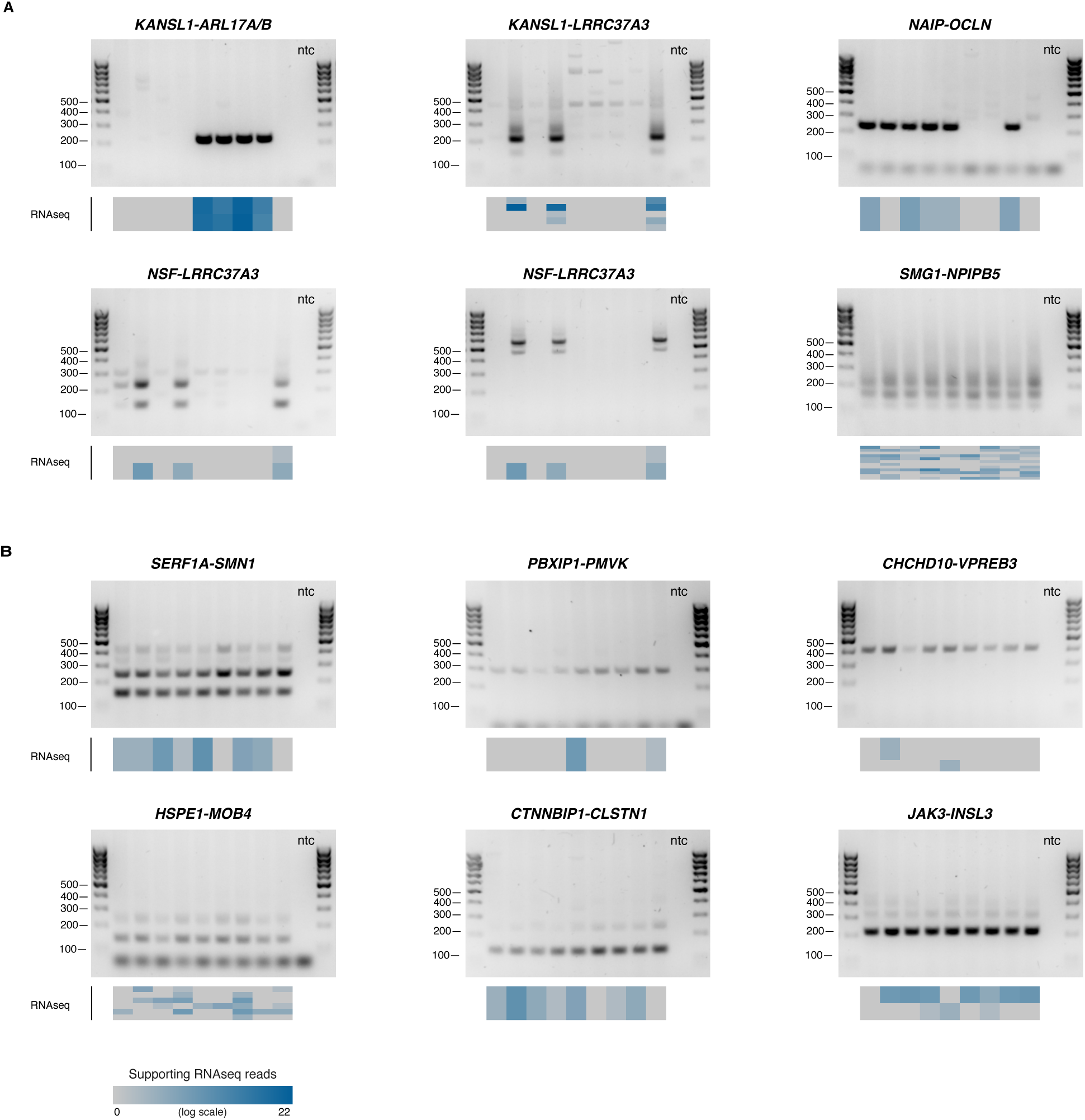
Fusion transcripts exhibit present/absent variation between individuals. **A-B:** Gel electrophoresis analysis of RT-PCR amplified fusion transcripts from 9 individuals, for genes in distant locations (**A**) and adjacent genes (**B**) in the reference genome. Heatmaps below each gel image show the expression level of each fusion transcript based on RNA-seq analysis of the same individual. Multiple boxes in the heatmap indicate different transcript variants of the same parental genes giving rise to the fusion transcript. Ntc: no template control.

### Variable fusion transcripts are located in highly duplicated regions of the genome associated with neurodevelopmental and neurodegenerative phenotypes

To elucidate the underlying genetics of fusion transcripts with clear inter-individual variations, we examined the location of fusion transcript junctions more closely. Among the fusions we examined by RT-PCR, those exhibiting presence/absence variations were located within genome regions 5q13.2 (**Figure 5**) and 17q21.31 (**Figure 6**). These loci are enriched with segmental duplications, and structural variations in these regions have been associated with neurodevelopmental and neuromuscular disorders^37–39^. Due to the high density of segmental duplications, these loci show considerable variation within the healthy human population. Notably, all of the variable fusion transcripts were frequently detected in the human population and not detected in RNA-seq data from chimpanzee or rhesus macaque (p<0.01; **Supplemental Figure 5**).

**Figure 5.**
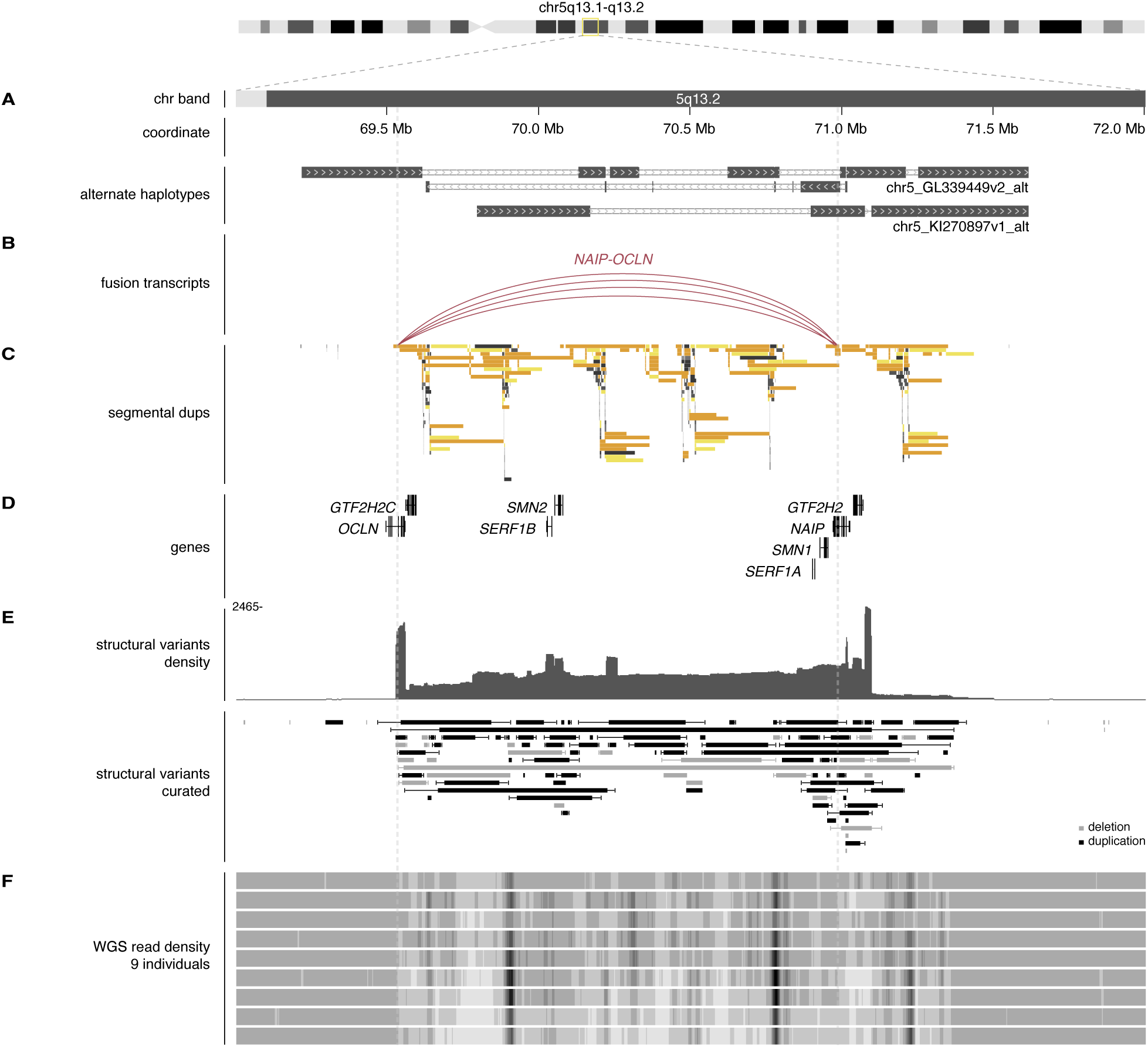
Variable fusion transcripts are located in the highly duplicated, copy-number variable locus 5q13.2, associated with Spinal Muscular atrophy (SMA). **A:** Chromosome band, hg38 coordinate and location of alternate haplotypes. **B**: Variable fusion transcript locations. Arcs connect the upstream and downstream fusion points of a single detected fusion transcript from short-read RNA-seq analysis. More than one arc joining the same two genes indicates multiple fusion transcripts were detected, fusing at different locations within the gene(s). **C**: Segmental duplication locations. **D**: Selected genes (GENCODE v44). **E**: Structural variants (deletions and duplications) found in healthy individuals from the Database of Genomic Variants (DGV). Both the density and a selection of curated variants found in multiple studies in the DGV are shown. **F**: Read density of WGS data from 9 healthy individuals (human iPSC lines) mapped to hg38, indicating copy number variability in the region.

**Figure 6.**
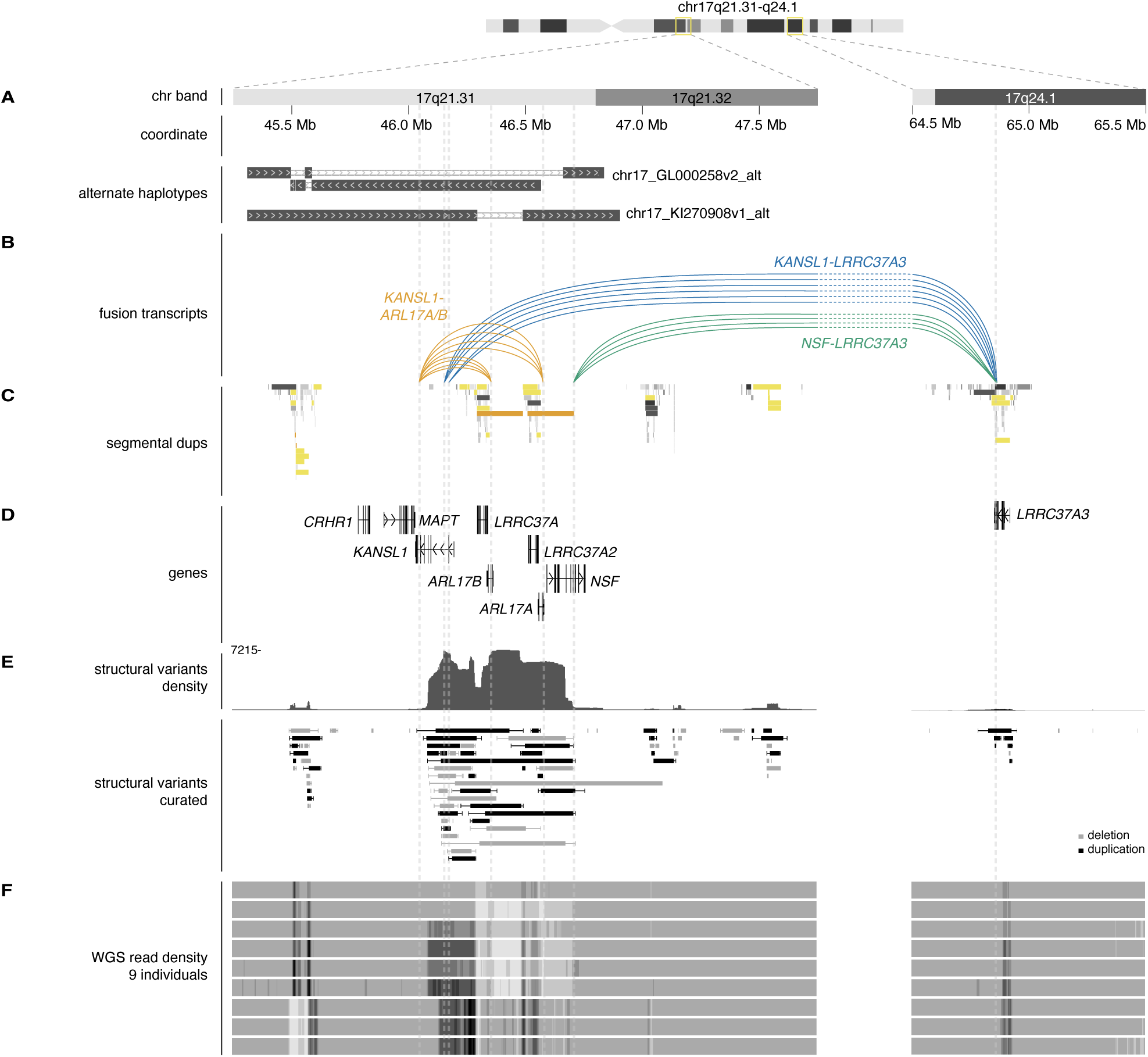
Variable fusion transcripts are located in the highly duplicated, copy-number variable locus 17q21.31, associated with neurodevelopmental and neurodegenerative disorders. **A**: Chromosome band, hg38 coordinate and location of alternate haplotypes. **B**: Variable fusion transcript locations. Arcs connect the upstream and downstream fusion points of a single detected fusion transcript from short-read RNA-seq analysis. More than one arc joining the same two genes indicates multiple fusion transcripts were detected, fusing at different locations within the gene(s). **C**: Segmental duplication locations. **D**: Selected genes (GENCODE v44). **E**: Structural variants (deletions and duplications) found in healthy individuals from the Database of Genomic Variants (DGV). Both the density and a selection of curated variants found in multiple studies in the DGV are shown. **F**: Read density of WGS data from 9 healthy individuals (human iPSC lines) mapped to hg38, indicating copy number variability in the region.

One of the variable fusion transcripts, *NAIP-OCLN*, originated from the 5q13.2 locus, which exhibits substantial genetic diversity and structural complexity (**Figure 5A-B**). Population studies have identified multiple distinct structural haplotypes and segmental duplications at 5q13.2 (**Figure 5A**; **C**). The region harbours the *SMN1* and *SMN2* genes (**Figure 5D**), with mutations or deletions in these genes leading to Spinal Muscular Atrophy (SMA), a severe neurodegenerative disorder marked by progressive muscle wasting and weakness^39–41^. While variations in *SMN1* or *SMN2* are the cause of the condition, SMA is also associated with variations in the copy number of *NAIP*^42–44^. The complex genomic architecture of 5q13.2 includes polymorphic inversions, deletions and duplications that create a dynamic landscape for gene expression regulation (**Figure 5E**). These variations in genome structure are common both within the healthy population (**Figure 5F**) and in SMA, where they can contribute to the diverse clinical presentations of the disease^45–47^. Additionally, CNVs in this region are associated with other conditions, such as susceptibility to infections and inflammatory responses, due to the involvement of the *NAIP* gene in immune pathways^48,49^.

The other variable fusion transcripts originated from genes in the 17q21.31 locus (**Figure 6A-B**), one of the most variable and complex regions of the human genome, which has also undergone dramatic reorganisation during recent human evolution^50,51^. We observed fusion events occurring between several genes in the locus, all originating from within segmental duplications (**Figure 6B-C**). 17q21.31 contains the *MAPT* gene, as well as *CRHR1*, *NSF*, *KANSL1*, and members of the *LRRC37* gene family, all of which have neural functions (**Figure 6D**)^52–60^. The region is known for its genomic instability, and frequent CNVs and complex rearrangements contribute to a high degree of inter-individual variability (**Figure 6E-F**). One specific recurrent copy number variation, 17q21.31 microdeletion, causes the neurodevelopmental disorder Koolen de Vries syndrome^37,38^. Eight structural sub-haplotypes exist within the normal human population^50,51^. Certain sub-haplotypes have been repeatedly associated with an increased risk of Parkinson’s disease (PD) and progressive supranuclear palsy (PSP) in genome-wide association studies (GWAS), while others are thought to confer a protective effect^61–64^ (**Supplemental Figure 6**). Having established the location of variable fusion transcripts within these highly duplicated, copy number-variable regions, we hypothesised that the expression of specific fusion transcripts is tightly linked to genomic haplotypes that exist for these loci.

### Fusion transcripts occur as a result of different genomic structural haplotypes

We next determined whether the presence or absence of variable fusion transcripts was indeed caused by different genomic structural arrangements in the human population. We focused on those fusion transcripts that exhibited clear presence/absence distinctions and were located in the 17q21.31 and 5q13.2 regions. We determined the full structure of *KANSL1-ARL17A/B*, *KANSL1-LRRC37A3*, *NSF-LRRC37A3* and *NAIP-OCLN* fusion transcripts using a combination of short-read RNA-seq, long-read Pacbio RNA-seq and RT-PCR of the full transcript structure followed by long-read Oxford Nanopore Technologies sequencing (**Figure 7A-C**; **Supplemental Figure 7**). Our analysis showed that the genes giving rise to these fusion transcripts were all located within segmental duplication regions, and were partial duplications of full-length genes (*KANSL1*, *NSF*, *NAIP* and *OCLN*) located elsewhere in the locus. The fusion transcript identified by FusionCatcher as *NAIP-OCLN* actually arose from the highly similar genes *NAIPP4* and *OCLNP1*. Similarly, *NSF-LRRC37A3* likely arose from the fusion of *NSFP1* and *LRRC37A2* genes and the *KANSL1* fusion genes originated from partially duplicated, unannotated copies of *KANSL1*, rather than the full-length annotated *KANSL1* gene.

**Figure 7.**
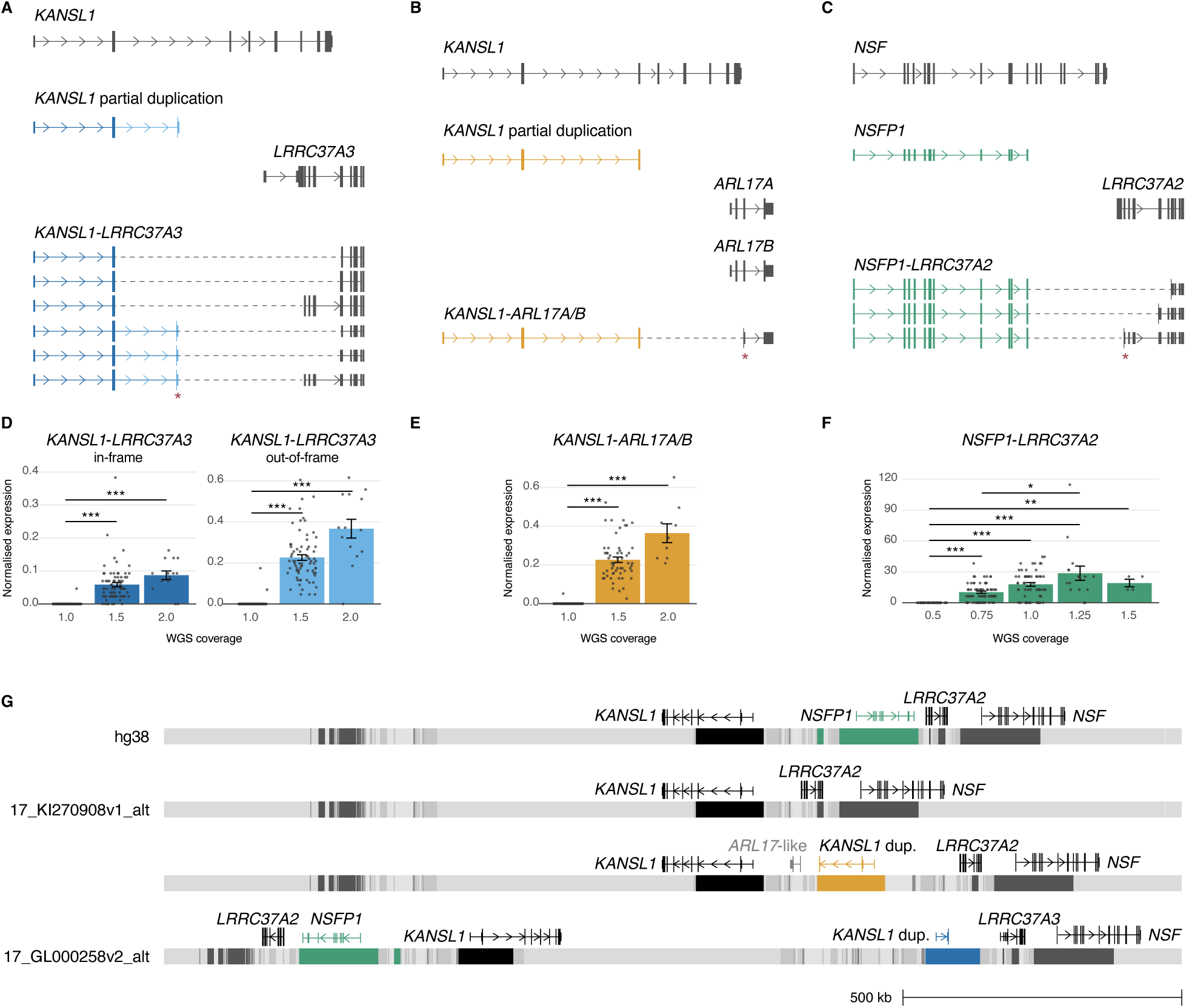
Variation in fusion transcript expression arises from different genomic structural haplotypes. **A-C**: Structure of *KANSL1-LRRC37A3*, *KANSL1-ARL17A/B* and *NSFP1-LRRC37A2* fusion transcripts, constructed based on Illumina, Pacbio and Oxford Nanopore Technologies sequencing data. Not all fusion transcript splice variants are depicted here; additional transcript variants are displayed in Supplemental Figure 7. **D-F**: Expression level of fusion transcripts based on genomic copy number of relevant segmental duplication (determined by WGS coverage). Fusion transcript expression is normalised relative to the non-duplicated parental control transcript expression level. Corresponding results for *NAIPP4-OCLNP1* and non-duplicated parental control transcripts are shown in Supplemental Figure 9. n=191 individuals; *p<0.05, **p<0.01, ***p<0.001. **G**: Structure of 17q21.31 genome region indicating proposed location of genes generating fusion transcripts. Coloured boxes represent segmental duplications containing partially duplicated genes *KANSL1* and *NSFP1*. Duplicated genes causing the fusion event are shown in colour. Organisation of all structural sub-haplotypes is displayed in Supplemental Figure 7.

To analyse if structural genetic variation was directly associated with the expression of fusion transcripts from the 17q21.31 and 5q13.2 loci, we analysed WGS data from 191 individuals from the Hipsci database. We quantified genome coverage in the region of the genes involved in fusion transcript formation, all located within segmental duplications. We found that for the region containing the *KANSL1* partial duplication, all individuals displayed a coverage ratio clustered around 1.0, 1.5 or 2.0, relative to a neighbouring single-copy genome region. This indicated inter-individual differences in the number of copies of this segmental duplication in the locus (**Supplemental Figure 8**). Similarly, individuals’ WGS coverage for the regions containing the *NSFP1* and *OCLNP1* duplications also clustered around discrete ratios relative to corresponding single-copy regions (**Supplemental Figure 8**). Using these WGS coverage ratios, we classified individuals into copy number groups and compared this genotype information to expression of the relevant fusion transcript in short-read RNA-seq data (**Figure 7D-F**; **Supplemental Figure 9**). Our analysis confirmed that the presence and level of fusion transcript expression was a direct result of variations in copy number at the corresponding genome region. Expression of the *KANSL1* fusion transcripts was completely absent or extremely rare in individuals with a copy number of 1.0, who presumably lack any segmental duplication of the *KANSL1* region (**Figure 7D-E**). Fusion transcript expression increased proportionally with genomic copy number in individuals with copy numbers of 1.5 or 2.0, indicative of the presence of a heterozygous or homozygous *KANSL1* duplication, respectively. Expression was also clearly linked to the genomic copy number group for the *NSFP1-LRRC37A2* and *NAIPP4-OCLNP1* fusion transcripts (**Figure 7F**; **Supplemental Figure 9**). We also examined the parental, full length genes which were the original source of the partially duplicated genes generating these fusion transcripts, and found no difference in expression between genotype groups (**Supplemental Figure 9**).

The pattern of fusion transcript expression we observed is consistent with the established structural sub-haplotypes of the 17q21.31 locus (**Figure 7G**; **Supplemental Figure 7**)^50,51^. Particular sub-haplotypes give rise to *KANSL1-ARL17A/B* and *KANSL1-LRRC37A3* due to the positioning of a partial *KANSL1* duplication upstream of *LRRC37A3* or an *ARL17*-like sequence, respectively. Fusion transcripts are initiated at the *KANSL1* partial duplication and continue via transcriptional readthrough into the downstream gene. Similarly, certain sub-haplotypes generate *NSFP1-LRRC37A2* fusions due to the positioning of a segmental duplication containing *NSFP1* upstream of *LRRC37A2*. We detected supporting Ribo-seq reads for *KANSL1-ARL17A/B* and *NSFP1-LRRC37A2* in one individual, indicating these fusions may be actively translated (**Supplemental Figure 10**). Both of these transcripts are fused out-of-frame and encode truncated versions of the full-length KANSL1 or NSF proteins. Notably, full-length in-frame *KANSL1-LRRC37A3* transcripts were present in Pacbio and Oxford Nanopore long-read sequencing data, and these transcripts would have the potential to produce a chimeric *KANSL1-LRRC37A3* protein (**Supplemental Figure 10**). The 17q21.31 sub-haplotype that generates *KANSL1-LRRC37A3* fusion transcripts has been repeatedly linked with decreased risk of multiple diseases, including PD and PSP^61–64^. While this warrants further investigation, the 17q21.31 locus is an example of presence or absence of a particular fusion transcript correlating almost perfectly with extremely well-established risk associations for several neurodegenerative diseases.

In summary, we demonstrated that many fusion events occur in the healthy human population, and a large number of these may have the potential to produce functional proteins. Some fusion transcripts exhibit pronounced inter-individual variation and are generated from human-specific segmental duplications that vary in copy number in the modern population. We elucidated how specific structural variants give rise to different fusion transcripts by characterising the association between sub-haplotypes in the 17q21.31 and 5q13.2 regions and the variability in fusion transcript formation, illustrating the genomic diversity that contributes to these patterns.

## Discussion

This study provides novel insights into the extent of fusion transcript expression in the human population and how variations in fusion expression can be driven by different genomic structural haplotypes. We identified hundreds of distinct fusion events occurring in non-cancerous human brain tissue. Most fusion transcripts were detected in only a few individuals, but some were very frequently expressed within the population. Our analysis revealed that certain fusion transcripts exhibited pronounced presence/absence differences in expression between individuals. All of these variable fusions were located in highly duplicated and unstable regions of our genome, associated with benign and pathogenic CNVs and harbouring genes involved in neurodevelopment and neurodegeneration. Additionally, we detected many fusion transcripts arising from transcriptional readthrough of nearby genes in single-copy genome regions that do not commonly exhibit structural variation. Unlike the variable fusion transcripts, these readthrough transcripts showed evidence of expression in most or all individuals, albeit at highly variable levels.

We further demonstrated that fusion transcripts with inter-individual differences in expression often arose from loci rich in human-specific segmental duplications. Most of these loci are genetically unstable and display a high level of copy number variation in the human population. The highly variable structure of these loci illustrates how segmental duplications can lead to the formation of novel fusion transcripts, providing a new source of transcriptomic diversity and evolutionary innovation in the human population. Intriguingly, the structural haplotype at 17q21.31 that gives rise to *KANSL1-LRRC37A3* fusion transcripts has been repeatedly associated with decreased risk of multiple diseases, including PD and PSP, in GWAS^61–65^. Conversely, a group of 17q21.31 sub-haplotypes, including the one which gives rise to the *KANSL1-ARL17A/B* fusion transcripts, have been repeatedly associated with an increased risk for these diseases. As such, the 17q21.31 locus represents a case where fusion transcripts segregate almost perfectly with GWAS risk haplotypes. While the causal variants in the 17q21.31 locus remain elusive, the fusion transcripts identified in this study should be considered as candidates underlying these differences in predisposition to 17q21.31-associated neurological disorders.

A significant proportion of the fusion transcripts we identified bioinformatically resulted from the fusion of two protein-coding regions, suggesting they may have protein-coding potential. Among these CDS-CDS fusions, the majority occurred in-frame, indicating their capacity to generate novel chimeric proteins. Typically, transcripts encoding out-of-frame CDS-CDS variants, as well as other fusion categories predicted to produce a frameshift, may be expected to be cleared by NMD. However, we observed some out-of-frame fusion transcripts expressed at high levels, including *KANSL1-ARL17A/B*, which also showed evidence of translation in Ribo-seq. This suggests that NMD does not always effectively downregulate out-of-frame transcripts. If such fusion transcripts persist, they could potentially produce truncated proteins with novel functions. Several examples support this concept: the truncated human-specific *NOTCH2NL* genes, derived from partial segmental duplication of *NOTCH2*, express stable transcripts and proteins even though they would typically be expected to undergo NMD^18^. Similarly, the human-specific genes *SRGAP2* and *ARHGAP11A* are truncated forms of their respective parental genes, yet they have evolved to perform essential functions as novel proteins^13,20^. What these truncated transcripts have in common with *KANSL1-ARL17A/B* is their high level of expression in human cells.

An unresolved question is whether fusion transcripts of interest, either in-frame or out-of-frame, are translated into stable proteins. In this study, we found evidence of active translation using Ribo-seq data. However, detecting a fusion transcript in ribosome profiling data does not confirm the stability of the translated fusion protein product, and the expression and stability of fusion protein isoforms require further validation. Validation could involve targeted mass spectrometry techniques such as multiple reaction monitoring to sensitively detect specific peptides spanning the fusion junction, as well as overexpression of tagged fusion transcripts. Such studies could confirm whether fusion transcripts are not only translated, but also result in stable and functional protein products. Although this goes beyond the scope of our current study, it is essential for uncovering the evolutionary impact of these fusion transcripts and understanding how variation in their expression could underlie certain individual-specific traits. Intriguingly, even non-protein coding fusion transcripts may have a physiological impact, such as the long non-coding chimeric RNA *SLC45A3-ELK4* which regulates cell proliferation by its transcript, not via a translated protein^66^.

Our findings reveal that fusion transcripts are common in non-cancerous tissues. Surprisingly, many of the fusion transcripts we detected in healthy human brain tissues and iPSCs were previously reported as cancer-associated transcripts or cancer biomarkers^66–70^. While a possible role for these assumed oncogenic fusion transcripts should not be ruled out, our data demonstrating high levels of expression in non-cancerous tissues should caution against their use as reliable cancer biomarkers. For example, *KANSL1-ARL17A/B* was identified as a “cancer predisposition gene” after being detected in RNA-seq data from 52% (14/27) of human glioblastoma samples and only 12% (2/17) of non-neoplastic brain tissues from different individuals^69^. In contrast, we detected this fusion transcript in 44% (122/276) of non-cancerous human brain tissue samples and 35% (67/191) of iPSC lines from healthy donors, which is close to the previously reported frequency in glioblastoma. Many studies of fusion gene expression in cancer have very small sample sizes, and a fusion transcript that occurs in healthy tissues could be detected in tumour samples more often than controls due to inter-individual genetic variation. Larger population studies and functional experiments are required to properly establish a particular fusion event as oncogenic.

In summary, our study demonstrates the existence of many previously unannotated fusion transcripts in human brain tissues, highlighting their presence/absence across different individuals or significant inter-individual variation in their expression. We reveal that these fusion transcripts often arise from recent human-specific genomic rearrangements, particularly in regions enriched with segmental duplications, contributing to substantial inter-individual variability. This phenomenon may represent an untapped source of evolutionary innovation and genetic diversity. Fusion transcripts can bring together exons from unrelated genes to generate chimeric proteins containing novel combinations of protein domains. While many genes in our genome originated from similar fusion events in our evolutionary past^28^, it is intriguing to consider that this process may continue in the human species today. Fusion events between adjacent genes, often resulting from transcriptional readthrough, provide evolutionary flexibility, enabling the creation of new proteins with potentially adaptive functions and the development of new gene structures.

We further demonstrate that genetically unstable loci, particularly those affected by waves of segmental duplication during primate evolution, are key sources of new fusion genes, transcripts, and proteins. Some of these fusions are present in only a subset of the human population. Perhaps not coincidentally, many of these segmental duplication-rich loci contain critical neuronal genes, with variations in their genomic organisation linked to various neurological disorders^31^. Although gene paralogs resulting from segmental duplications are frequently annotated as pseudogenes, our findings show that they can be expressed as part of fusion transcripts, which may produce functional proteins. Thus, our study underscores the importance of investigating pseudogenes in segmental duplication-rich regions to assess their expression and protein-coding potential as components of the fusion transcripts identified here. This is essential because, even when in cases where a causal factor for a disease-associated locus is known, the penetrance and phenotypic expression of the associated disorders often varies widely among affected individuals. This suggests that other genetic elements within such loci likely contribute to the onset, severity, or progression of the disorders. We propose that fusion transcripts and the proteins they encode are promising candidates for explaining part of this inter-individual variation, as they may play critical roles in both health and disease, influencing physiological processes and disease susceptibility.

## Author contributions

Moses C., Vanini A., and Bocchi M.C. contributed to conceptualization, data curation, formal analysis, investigation, methodology, and validation. Wagner L. and van Loenen I. assisted with data curation, formal analysis, investigation, and methodology. Latour B.L. contributed to conceptualization, investigation, methodology, and project administration. Nadif Kasri N. and Jacobs F.M.J. provided conceptualization, funding acquisition, project administration, and supervision. Moses C., Vanini A., and Jacobs F.M.J. wrote the original draft, with all authors participating in the review and editing process. Jacobs F.M.J. is the corresponding author.

## Acknowledgements

We would like to thank Sabine Spijker, Guus Smit, Frank Koopmans, and Femke Roig Kuhn for their valuable consultation on fusion peptide detection in mass spectrometry data. We are also grateful to Wim de Leeuw for assistance with the compute cluster. This work was funded by an NWO ENW grant to F. Jacobs and N. N. Kasri. C. Moses was supported by an EMBO Postdoctoral Fellowship.

