## Supplemental methods for "Evolutionary hotspots of structural variation drive inter-individual differences in the expression of fusion transcripts in the human brain"

#### Samples and data sources

Human RNA-seq data for the genome-wide fusion transcript analysis were obtained from the Mayo RNA-seq Study (University of Florida, Mayo Clinic, Institute for Systems Biology, Emory University). Postmortem tissue from the temporal cortex of 276 North American Caucasian subjects was analysed. Data were accessed via Synapse (synapse.org; SynID syn8612203). To validate fusion transcripts using PacBio long-read sequencing, data from 6 individuals were accessed from three publicly available datasets[1,2] (downloads.pacbcloud.com/public/dataset/Kinnex-full-length-RNA/). Available data from 3 healthy donors and 3 unrelated patients with familial Parkinson's disease were used for ribosome profiling analysis[3]. RNA-seq data from chimpanzee and rhesus macaque brain (NCBI BioProjects PRJEB33938, PRJNA793060, PRJNA271912, PRJNA285021, PRJNA393104, PRJNA236446) were analysed to compare fusion transcript expression between human and other primate species. To compare fusion transcript expression with structural variant haplotypes in the human population, RNA-seq and WGS data from the Human Induced Pluripotent Stem Cell Initiative[4,5] (HipSci) were accessed via the European Nucleotide Archive (ENA). Human iPSC lines from HipSci were accessed via the European Bank for induced pluripotent Stem Cells (EBiSC) and used for RT-PCR validation of fusion transcripts.

#### Short-read fusion transcript analysis

Fusion transcript detection in Illumina short-read RNA-seq data was performed using FusionCatcher[6]. FusionCatcher identifies reads and read pairs that map onto more than one annotated gene using several aligners, including Bowtie, BLAT, and STAR. FusionCatcher (version 1.33) was installed via conda and used with the in-built human reference genome hg38. FusionCatcher typically flags known fusions found in normal tissue databases; however, we did not remove these from our results. We performed additional filtering of the FusionCatcher results, excluding those flagged as similar reads, duplicates, short repeats, long repeats, pairs of pseudogenes, paralogs, ambiguous, and those with a high number of common mapping reads between parental genes, indicating sequence similarity. We also excluded fusions where the junction overlapped with the hg38 RepeatMasker track.

The combined list of all fusion transcripts discovered in all individuals from the Mayo human RNA-seq dataset is provided in Supplemental File 2. We only considered fusion transcripts detected in two or more individuals in FusionCatcher analysis. The fusion junction sequences from each transcript, consisting of 43 nucleotides immediately upstream and 43 nucleotides downstream of the junction splice site, were used to construct a custom reference (provided as Supplemental File 3). Fastq reads were aligned against this custom reference using bowtie2 to discover additional fusion-supporting reads not identified by standard FusionCatcher analysis. Score settings for bowtie2 alignment are provided in Supplemental File 4. Data from FusionCatcher and bowtie2 alignment were further analysed in R, using karyoplotR and circlize packages to visualise intrachromosomal and interchromosomal fusion locations, respectively. We compared which fusion transcripts were located within segmental duplications (SD locations were obtained from the Duplications of >1000 Bases of Non-RepeatMasked Sequence track via the UCSC Genome Browser[7,8]). All scripts used throughout the study are available at Code Ocean.

#### Long-read fusion transcript analysis

Of the 3 PacBio long-read sequencing datasets used for validation of fusions from short-read sequencing data, the first (downloads.pacbcloud.com/public/dataset/Kinnex-full-length-RNA/) and second[1] were used to validate fusion transcripts found in short-read RNA-seq data, while the third[2] was used solely to validate the full-length structure of *KANSL1-ARL17A/B*, *KANSL1-LRRC37A3*, *NAIPP4-OCLNP1*, and *NSFP1-LRRC37A2* fusion transcripts. Analyses were performed on full-length, non-concatemer (FLNC) reads. For the PacBio Kinnex full-length RNA dataset (downloads.pacbcloud.com/public/dataset/Kinnex-full-length-RNA/), FLNC reads were available directly. For the second dataset[1], FLNC reads were generated from CCS reads using the Iso-Seq pipeline (https://isoseq.how/). Briefly, HiFi reads (predicted accuracy ≥Q20) were first extracted from the CCS reads of each sample using extracthifi, primers were removed using lima, and isoseq refine was used for trimming PolyA tails and concatemer removal. For the third dataset[2], HiFi reads were generated from subreads.bam files using the ccs tool (https://github.com/PacificBiosciences/ccs), and FLNC reads were then generated using lima and isoseq refine Iso-Seq tools as described for the second dataset. After all FLNC files were obtained, minimap2 was used to index and map the FLNC long reads onto the 717 fusion transcript junction sequences. For mapping, a maximum of 3 mismatches were allowed in each fusion junction sequence of 86 nucleotides.

De novo fusion transcript detection in Iso-Seq HiFi data was subsequently performed on the samples of the first and second dataset using pbfusion (github.com/pacificBiosciences/pbfusion/). Iso-Seq HiFi reads (FLNC) were first aligned to the human genome (GRCh38.p13.genome.fa) with pbmm2 (github.com/PacificBiosciences/pbmm2). Then pbfusion was run on aligned reads, allowing the emission of both medium and low quality fusions. Full results of fusion transcripts identified in the de novo pbfusion analysis and the validation of short-read data using minimap2 are provided in Supplemental File 5.

**Protein-coding potential of fusion transcripts**

To visualise the *in silico* protein-coding potential of the fusion transcript dataset (as predicted by FusionCatcher), we used the Python package Matplotlib. The fusion junctions of upstream and downstream fusion genes were classified as falling within the UTR (untranslated region), CDS (CoDing Sequence), exonic (no-known-CDS), or intronic. Fusion transcripts where both upstream and downstream fusion junctions fell within a CDS were further classified as in-frame or out-of-frame. All categories are based on the coding potential and reading frame of the annotated parental genes; however, it is possible that fusion transcripts may contain novel open reading frames that are not present in the parental genes, even when one or both of the parental genes are non-coding. Seven fusion transcripts were excluded from visualisation as FusionCatcher was unable to classify the protein-coding potential of their sequences. Conserved domains in the predicted protein structures were identified using the NCBI conserved domains search tool (ncbi.nlm.nih.gov/Structure/cdd/wrpsb.cgi).

To collect evidence for the translation of fusion transcripts, we analysed Ribo-seq data from 3 healthy donors and 3 unrelated patients with familial Parkinson's disease[3]. SRA files (GEO accession numbers GSM2402496, GSM2402497, GSM2402498, GSM2402499, GSM2402500, GSM2402501) were downloaded and converted to fastQ format using the SRA Toolkit packages prefetch and fastq-dump[9], followed by trimming with Trim Galore! (www.bioinformatics.babraham.ac.uk/projects/trim_galore/) and quality control checks using FastQC (www.bioinformatics.babraham.ac.uk/projects/fastqc/). Only reads longer than 16 bp were retained. The same custom reference used for bowtie2 alignment of RNA-seq reads, consisting of 43 nucleotides immediately upstream and 43 nucleotides downstream of the junction splice site (Supplemental File 3), was used to align Ribo-seq reads mapping to the fusion junctions. Bowtie2[10] was used to index the fusion transcript sequences and map the Ribo-seq reads onto the custom reference. Zero mismatches were allowed within a 16 bp seeding sequence, and only Ribo-seq reads spanning at least 5 bp upstream and 5 bp downstream of the fusion junction were retained. To exclude Ribo-seq reads that might map to multiple different transcripts, a BlastN[11] analysis was performed against the transcriptome (GENCODE v44). Ribo-seq reads that matched other known transcripts with 100% coverage were discarded. The set of control transcript junctions was generated from all exon junctions in GENCODE v44 (including only those where the exonic sequence on each side of the junction extended for at least 43 nucleotides). Three sets of 717 control exon junction-spanning sequences were randomly selected and analysed in the same way as the 717 fusion transcript junctions (control junction sequences are provided in Supplemental File 3). For the control set, Ribo-seq reads that matched known transcripts (other than the intended transcript) with 100% coverage were discarded.

**Cross-species fusion transcript analysis**

Fastq files from chimpanzee and rhesus macaque (NCBI BioProjects PRJEB33938, PRJNA793060, PRJNA271912, PRJNA285021, PRJNA393104, PRJNA236446) were aligned using bowtie2 against the custom reference of fusion junctions detected by FusionCatcher in the human Mayo RNA-seq dataset (Supplemental File 3). Score settings were adjusted to allow for a greater number of mismatches based on the known substitution rate between humans and each species (Supplemental File 5). A list of the fusion transcripts detected only in humans at levels $p<0.05$ and $p<0.01$ is provided in Supplemental File 6.

### Locus diagrams

Data shown in the locus schematics for chr5q13.2 and chr17q21.31 were obtained via the UCSC Genome Browser[7,8,12]. WGS data were obtained from HipSci[4] (hipsci.org) and mapped to hg38 using BWA-MEM. SAM files were converted to BAM, sorted and indexed using samtools. BigWig coverage files were generated using bamCoverage and visualised against the UCSC Genome Browser, genome version hg38.

### RT-PCR validation of fusion transcripts

RT-PCR was conducted in 9 HipSci iPSC lines generated from different individuals[4,5]. PCR primers were designed in the exons up- and downstream of the fusion junction using NCBI Primer Blast (ncbi.nlm.nih.gov/tools/primer-blast/) to validate that primers did not amplify other known transcripts in the Human Nucleotide Set. RNA was extracted by phenol-chloroform extraction using TRIzol Reagent (Invitrogen). DNase treatment was performed using DNase I (Roche) and samples were cleaned up using the RNA Clean & Concentrator-5 Kit (Zymo). Two micrograms of total RNA was converted to cDNA using SuperScript II Reverse Transcriptase (Thermo Fisher Scientific) according to the First-Strand cDNA Synthesis Using SuperScript II RT protocol (Pub. No. MAN0001342). PCR was carried out using Phusion High-Fidelity DNA Polymerase (New England Biolabs) with GC buffer and standard reaction conditions. Products were visualised on agarose gel alongside MassRuler Low Range DNA ladder (Thermo Fisher Scientific). Primer sequences and annealing temperatures are displayed in Supplemental File 7.

### Variable fusion transcript expression and structural variant genotyping

Publicly available RNA-seq and WGS data from iPSC lines from the HipSci project[4,5] were used to establish structural variants causing variable fusion transcript expression. Bowtie2 mapping of fastq files against fusion junction sequences (43 nucleotides either side of the splice site, Supplemental File 3) was used to establish the number of supporting reads for variable fusion transcripts. Reference sequences of the corresponding full-length parental genes at the same exon junction were generated for each fusion transcript as controls for normalisation (Supplemental File 3). HipSci WGS data (already mapped to GRCh37/hg19) were used to establish each individual's copy number in structurally variable regions. Relevant regions of aligned reads in hg19 were retrieved using samtools view. The coverage of copy number variable regions relative to nearby single copy regions was quantified using bedtools multicov. For distinguishing the 17q21.31 inversion, the H1/H2 discriminating SNP rs8070723 was used, which is in linkage disequilibrium with the H1 orientation[13,14]. The frequency of this SNP in each individual was determined using bcftools mpileup at the coordinate chr17:44081064 (hg19).

### Variable fusion transcript isoform discovery

Isoform structure of variable fusion transcripts was determined using a combination of Pacbio and Oxford Nanopore Technologies (ONT) long-read sequencing. Pacbio sequencing data were analysed as described in the section above and reads supporting *KANSL1-ARL17A/B*, *KANSL1-LRRC37A3*, *NSFP1-LRRC37A2*, and *NAIPP4-OCLNP1* fusion transcripts were saved to fasta files. Fasta files for each fusion transcript were used to identify transcript isoforms using Isoquant (https://www.nature.com/articles/s41587-022-01565-y) with human reference genome hg38. Isoform output gtf files were visualised on the UCSC Genome Browser and used to construct transcript diagrams in Adobe Illustrator. To further experimentally validate the presence of full-length *KANSL1-LRRC37A3* fusion transcripts and their associated splice variants, RT-PCR was conducted to amplify the whole length of the fusion transcript using Phusion High-Fidelity DNA Polymerase (New England Biolabs). Primer sequences and annealing temperatures are displayed in Supplemental File 7. PCR products were run on agarose gel and purified using the QIAEX II Gel Extraction Kit (QIAGEN). Purified products were quantified following the Qubit dsDNA BR Assay Kit (Pub. No. MAN000232) or the Qubit 1X dsDNA HS Assay Kits (Pub. No. MAN0017455). ONT libraries were generated according to the protocol "Ligation sequencing amplicons - native barcoding (SQK-LSK109 with EXP-NBD104 and EXP-NBD114)," with the EXP-NBD103 kit used for barcoding. Sequencing was performed on the MinION (FLO-MIN106R9). Transcript isoform discovery was performed using Isoquant (github.com/ablab/IsoQuant) based on reference genome hg38.

**Data analysis and availability**

Analysis was performed on the University of Amsterdam Faculty of Science compute cluster. Associated code is available at Code Ocean. Scripts were written and executed in RStudio version 2022.02.2 and Bash version 4.2.46(2)-release. Bash jobs were executed by workload manager Slurm version 18.08.8. Raw data are available from the original sources. Processed data are available on request to the authors.
