## Supplemental figures for "Evolutionary hotspots of structural variation drive inter-individual differences in the expression of fusion transcripts in the human brain"

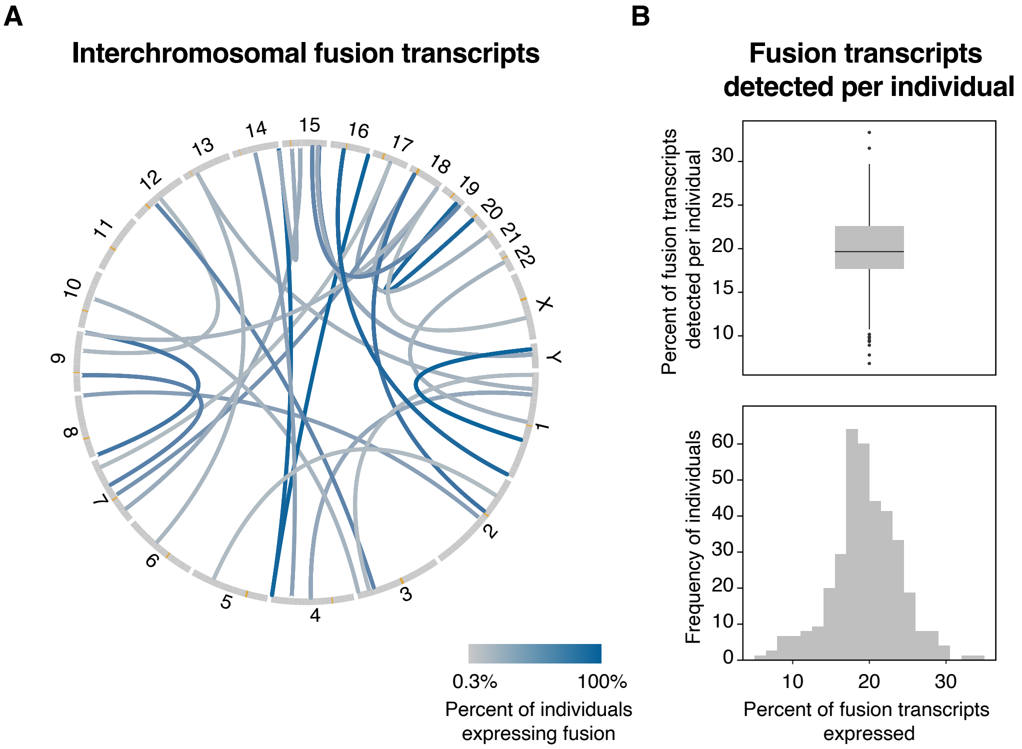


**Supplemental Figure 1. Extra features of genome wide fusion detection. A**: Representation of interchromosomal fusion transcripts detected in two or more individuals in postmortem human brain RNA-seq data from 276 individuals. Each blue arc connects the upstream and downstream fusion points of a single detected fusion transcript. The depth of colour of each arc corresponds to the percentage of individuals in which the fusion transcript was detected. Centromeres are depicted in orange. **B**: Percentage of the 717 fusion transcripts detected per individual.

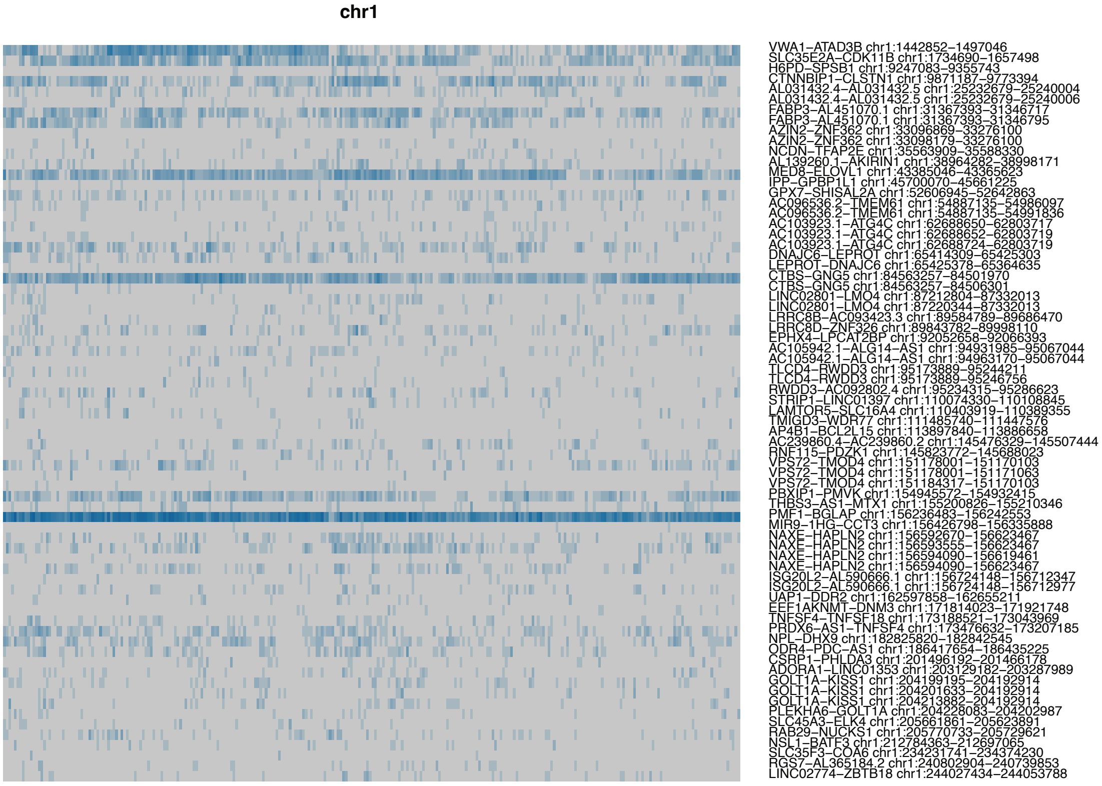


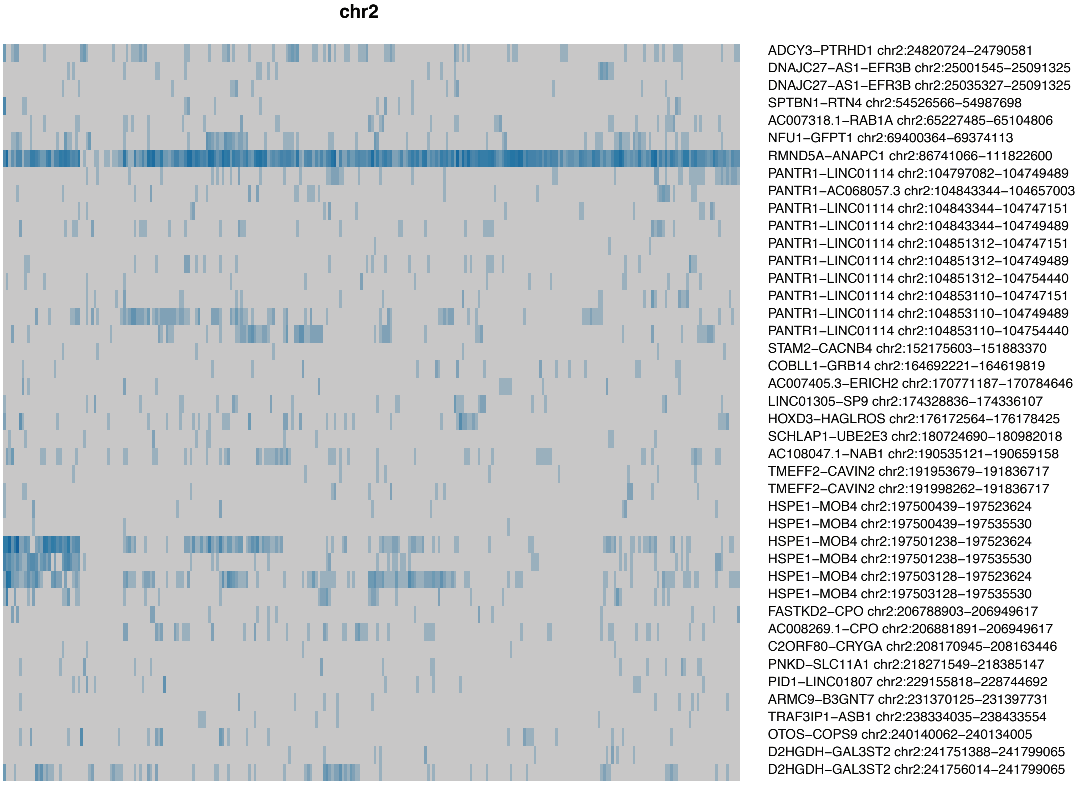


**Supplemental Figure 2. Fusion transcripts detected in short-read RNA sequencing data (all chromosomes).** Heatmaps displaying fusion transcripts detected per individual on each chromosome. Rows represent different fusion transcripts, and columns represent individuals. Depth of colour is proportional to the number of RNAseq reads supporting the fusion.

(continued next page)

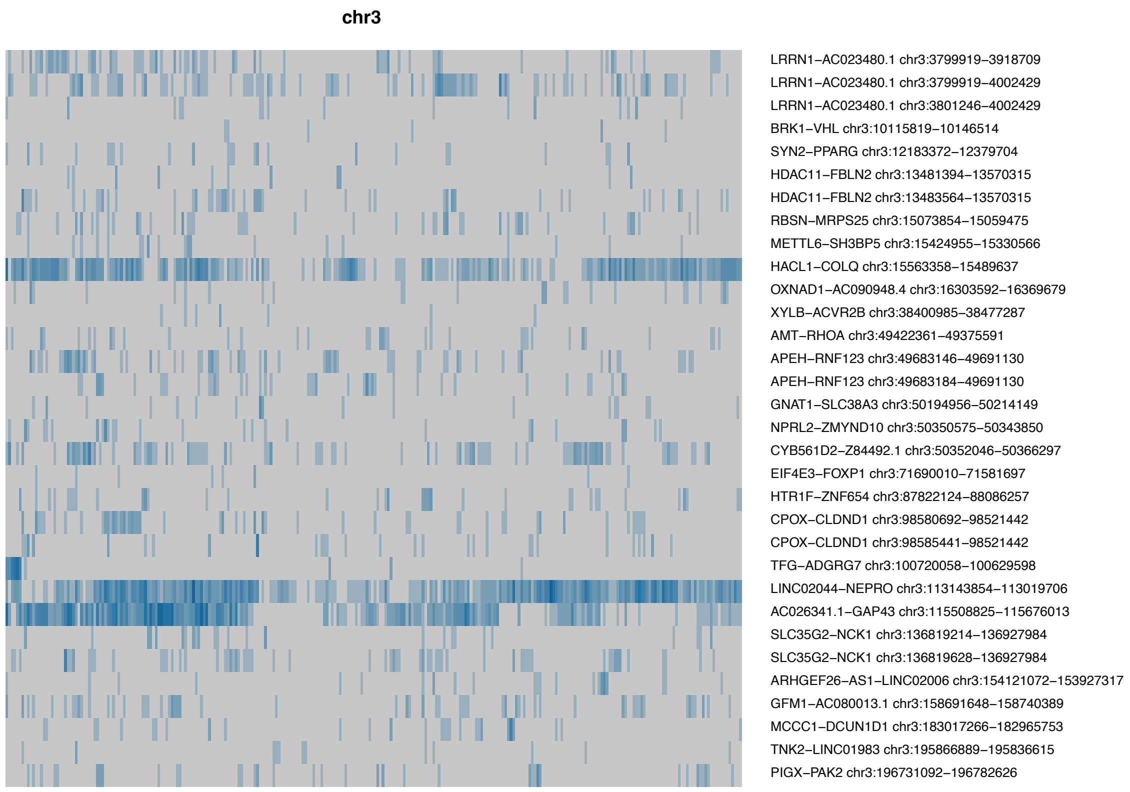


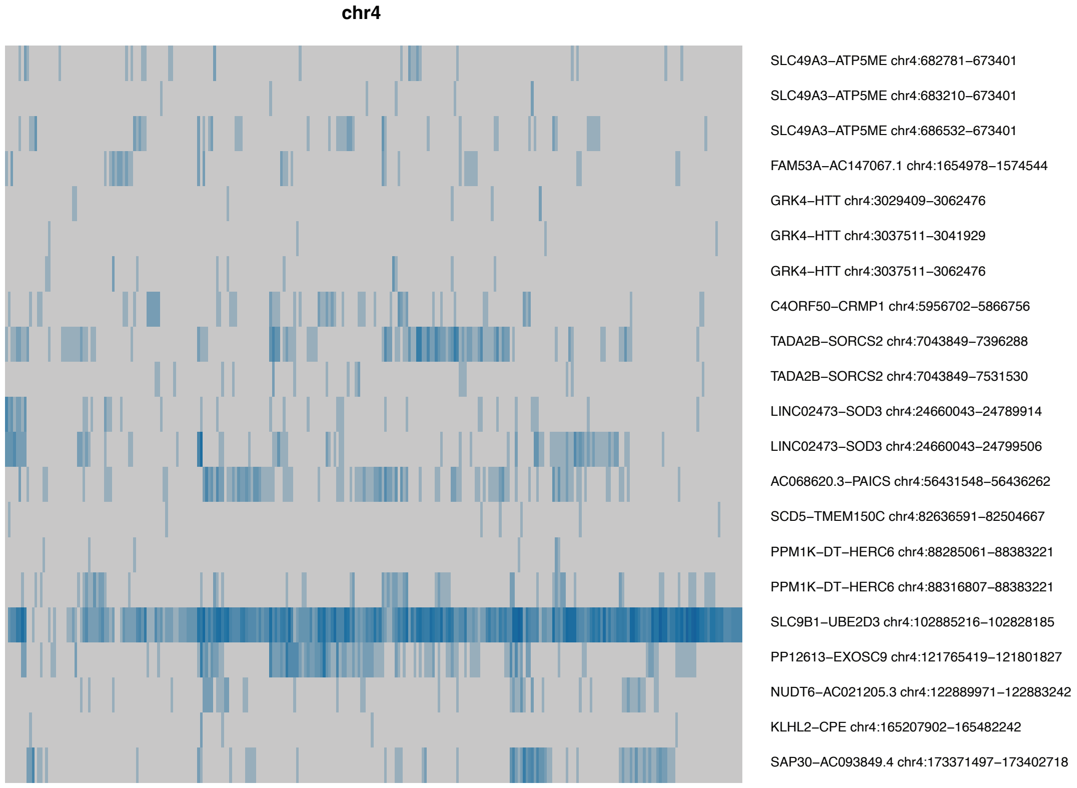


**Supplemental Figure 2. Fusion transcripts detected in short-read RNA sequencing data (all chromosomes).** Heatmaps displaying fusion transcripts detected per individual on each chromosome. Rows represent different fusion transcripts, and columns represent individuals. Depth of colour is proportional to the number of RNAseq reads supporting the fusion.

(continued next page)

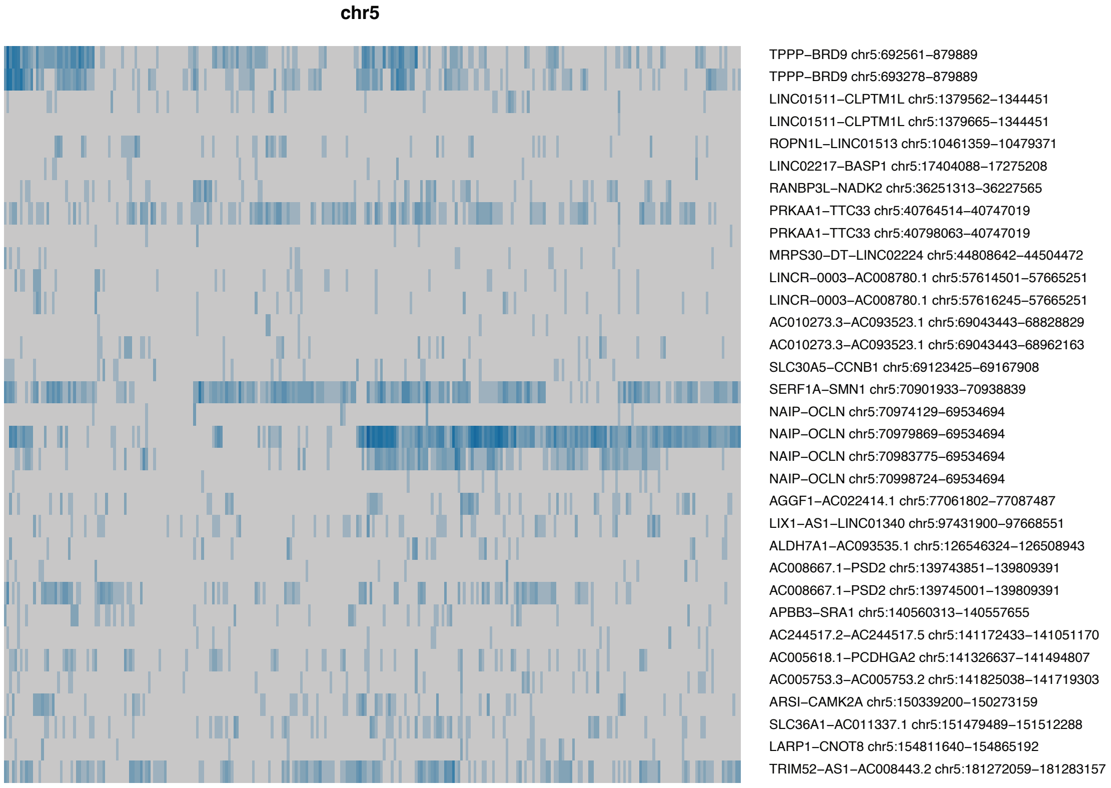


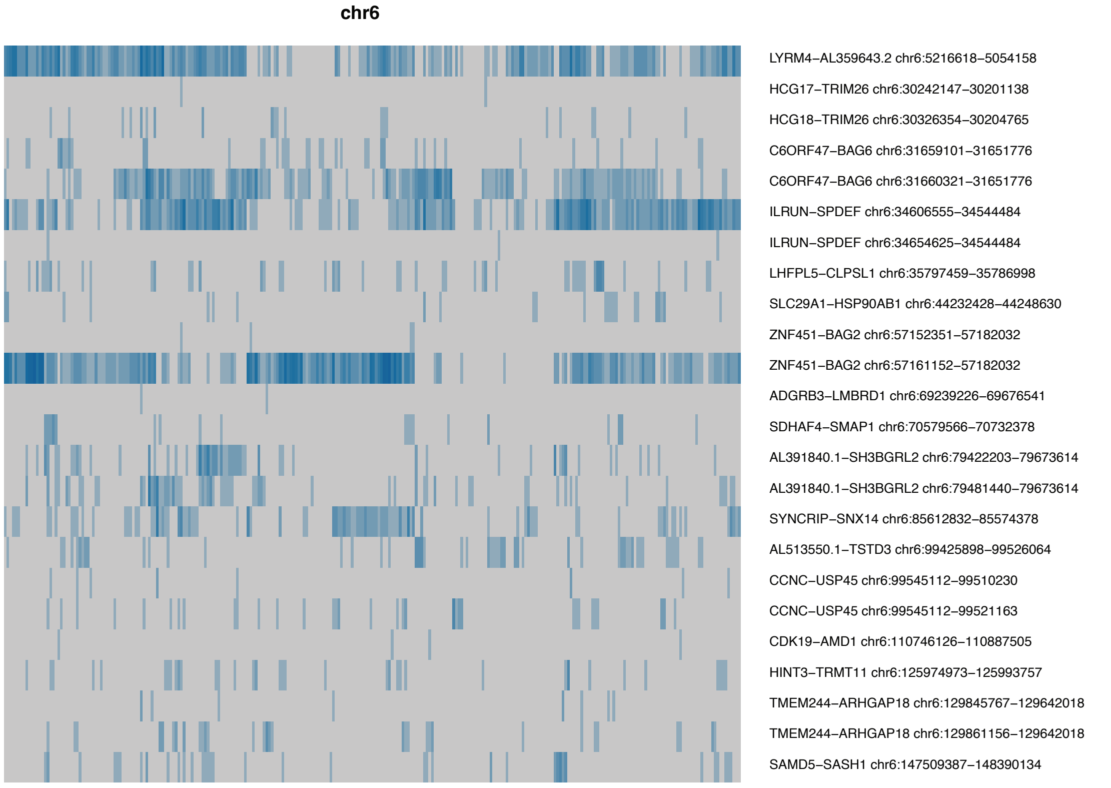


**Supplemental Figure 2. Fusion transcripts detected in short-read RNA sequencing data (all chromosomes).** Heatmaps displaying fusion transcripts detected per individual on each chromosome. Rows represent different fusion transcripts, and columns represent individuals. Depth of colour is proportional to the number of RNAseq reads supporting the fusion.

(continued next page)

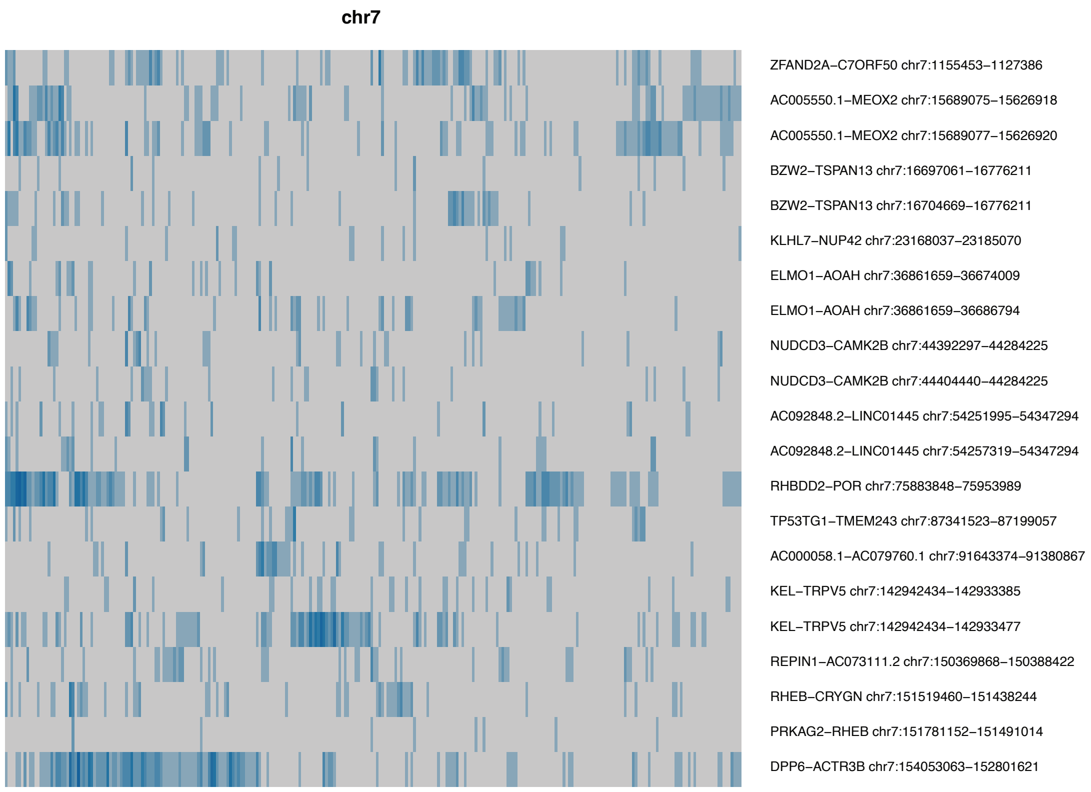


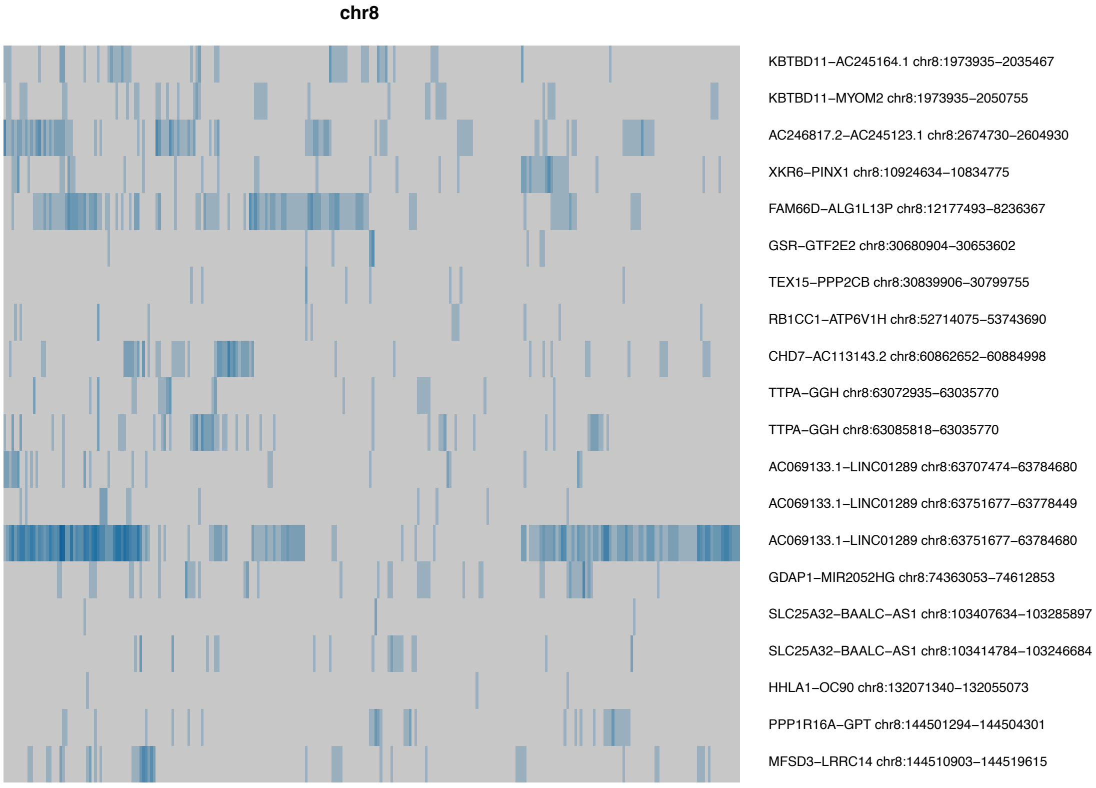


**Supplemental Figure 2. Fusion transcripts detected in short-read RNA sequencing data (all chromosomes).** Heatmaps displaying fusion transcripts detected per individual on each chromosome. Rows represent different fusion transcripts, and columns represent individuals. Depth of colour is proportional to the number of RNAseq reads supporting the fusion.

(continued next page)

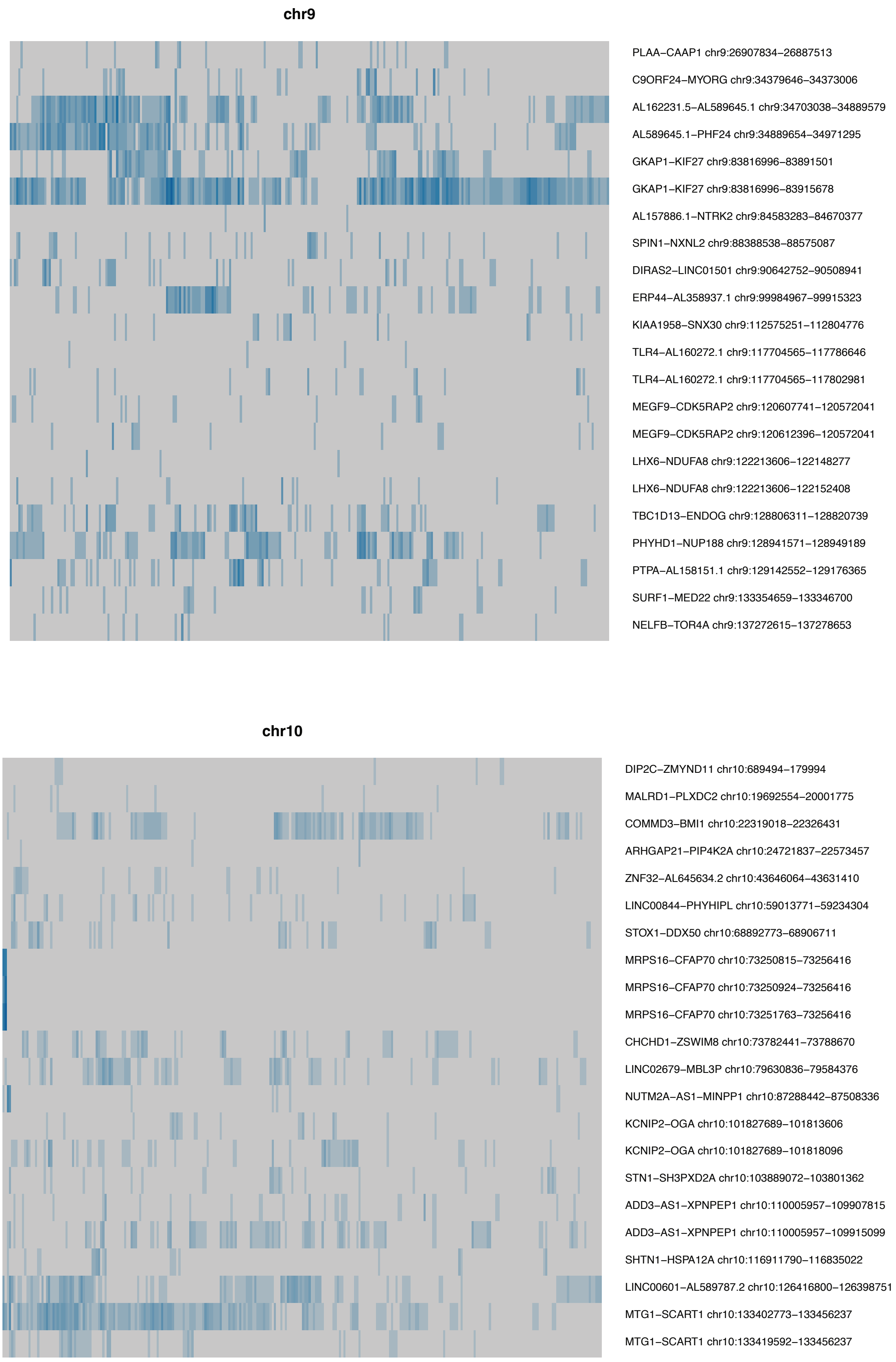


**Supplemental Figure 2. Fusion transcripts detected in short-read RNA sequencing data (all chromosomes).** Heatmaps displaying fusion transcripts detected per individual on each chromosome. Rows represent different fusion transcripts, and columns represent individuals. Depth of colour is proportional to the number of RNAseq reads supporting the fusion.

(continued next page)

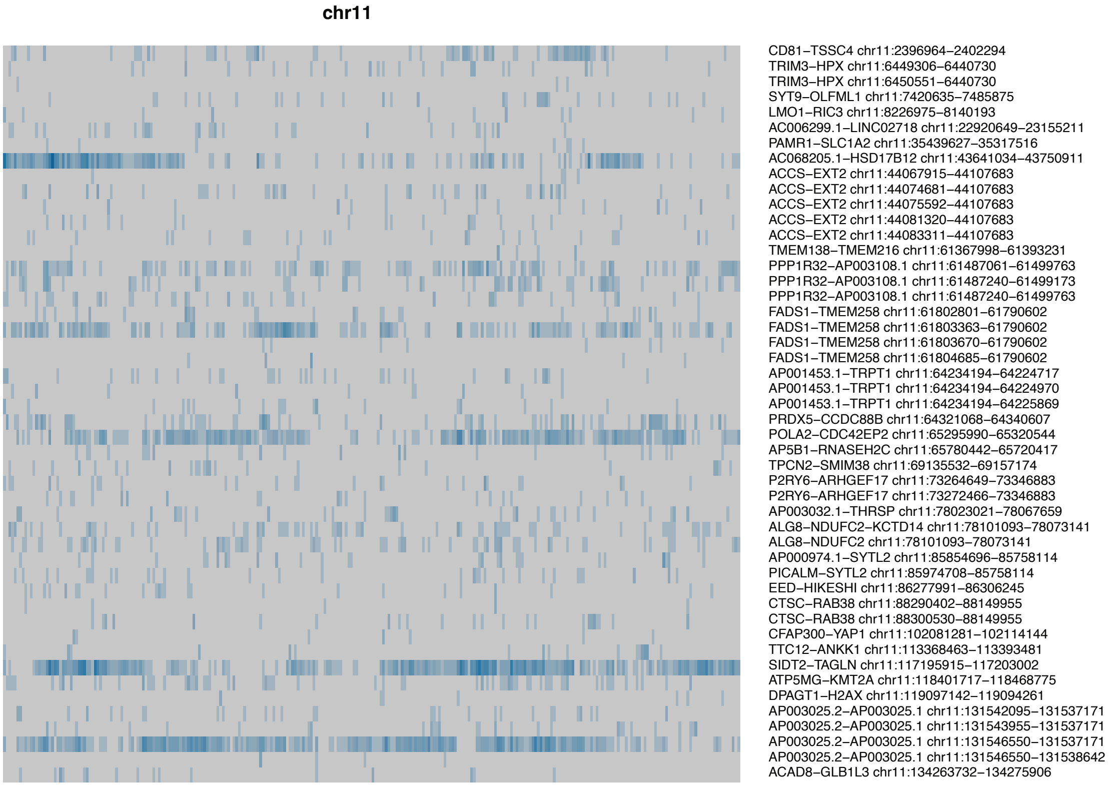


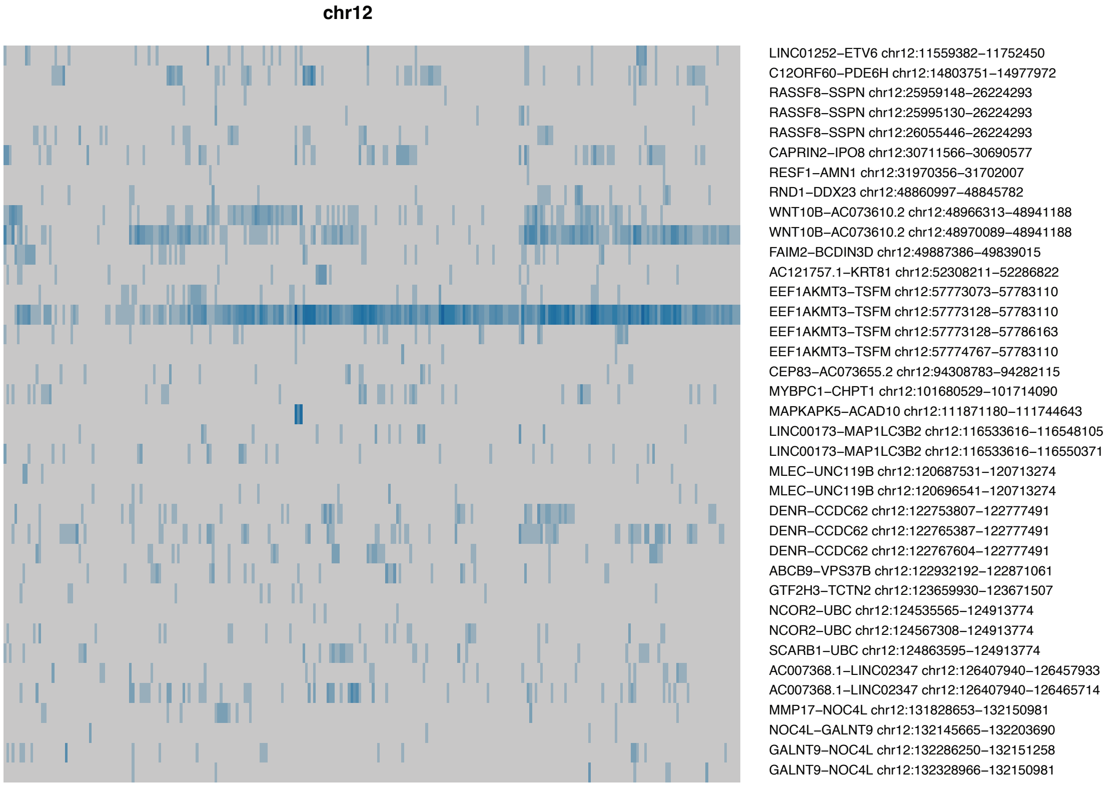


**Supplemental Figure 2. Fusion transcripts detected in short-read RNA sequencing data (all chromosomes).** Heatmaps displaying fusion transcripts detected per individual on each chromosome. Rows represent different fusion transcripts, and columns represent individuals. Depth of colour is proportional to the number of RNAseq reads supporting the fusion.

(continued next page)

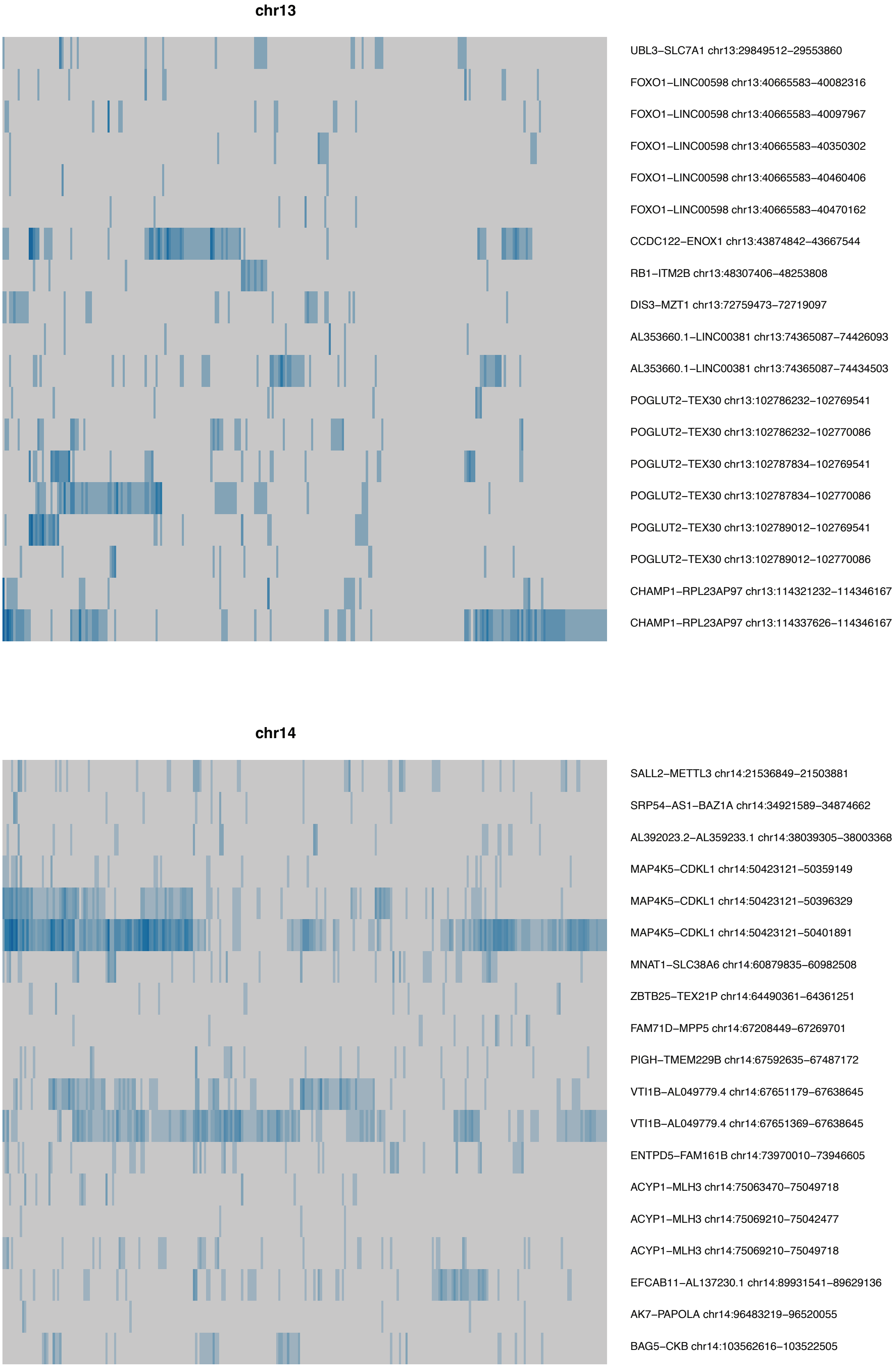


**Supplemental Figure 2. Fusion transcripts detected in short-read RNA sequencing data (all chromosomes).** Heatmaps displaying fusion transcripts detected per individual on each chromosome. Rows represent different fusion transcripts, and columns represent individuals. Depth of colour is proportional to the number of RNAseq reads supporting the fusion.

(continued next page)

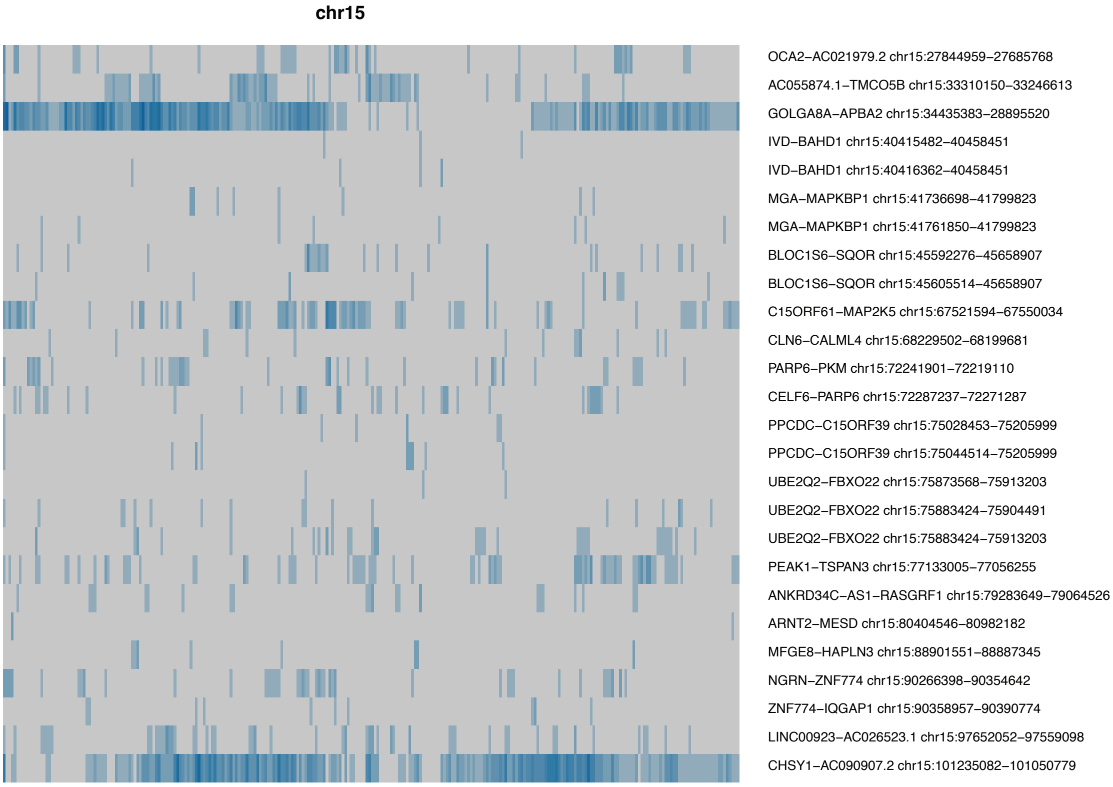


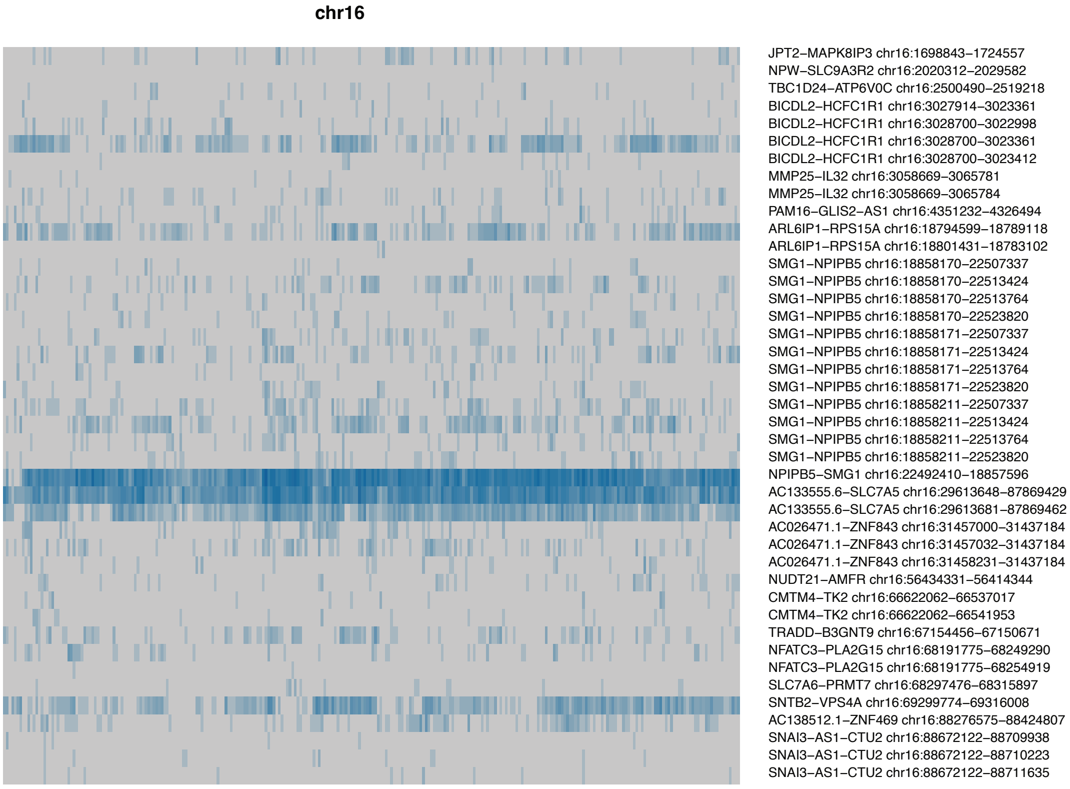


**Supplemental Figure 2. Fusion transcripts detected in short-read RNA sequencing data (all chromosomes).** Heatmaps displaying fusion transcripts detected per individual on each chromosome. Rows represent different fusion transcripts, and columns represent individuals. Depth of colour is proportional to the number of RNAseq reads supporting the fusion.

(continued next page)

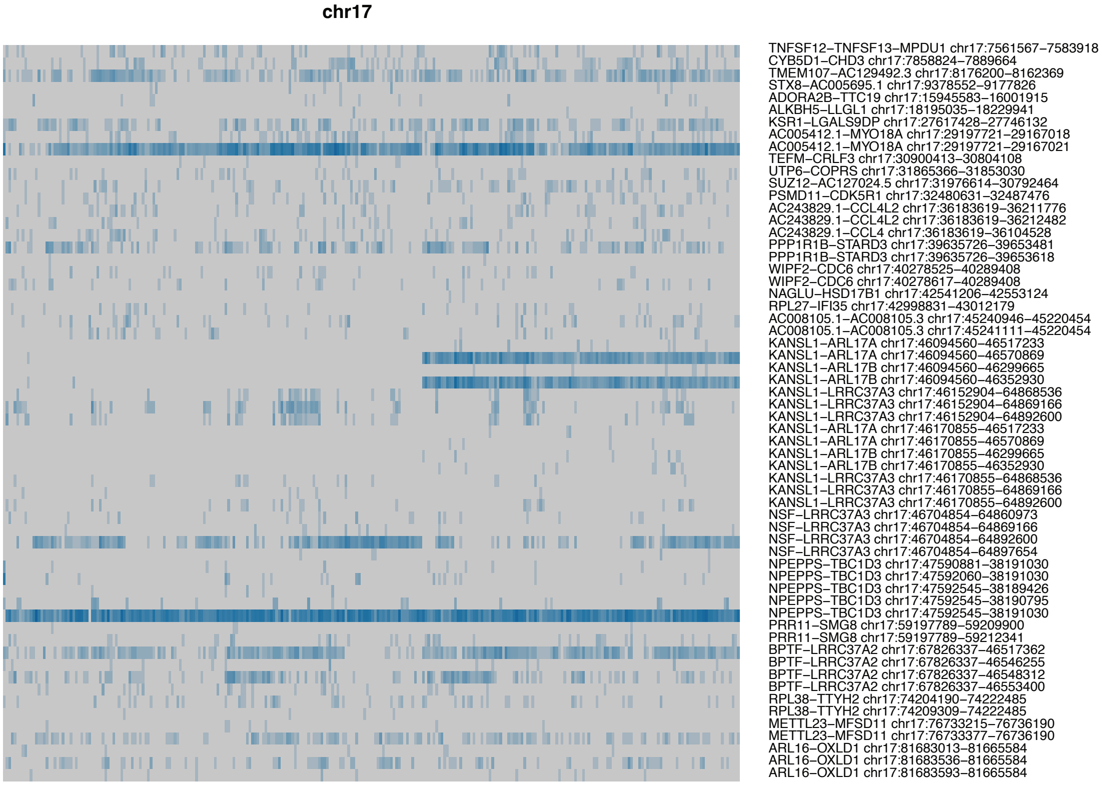


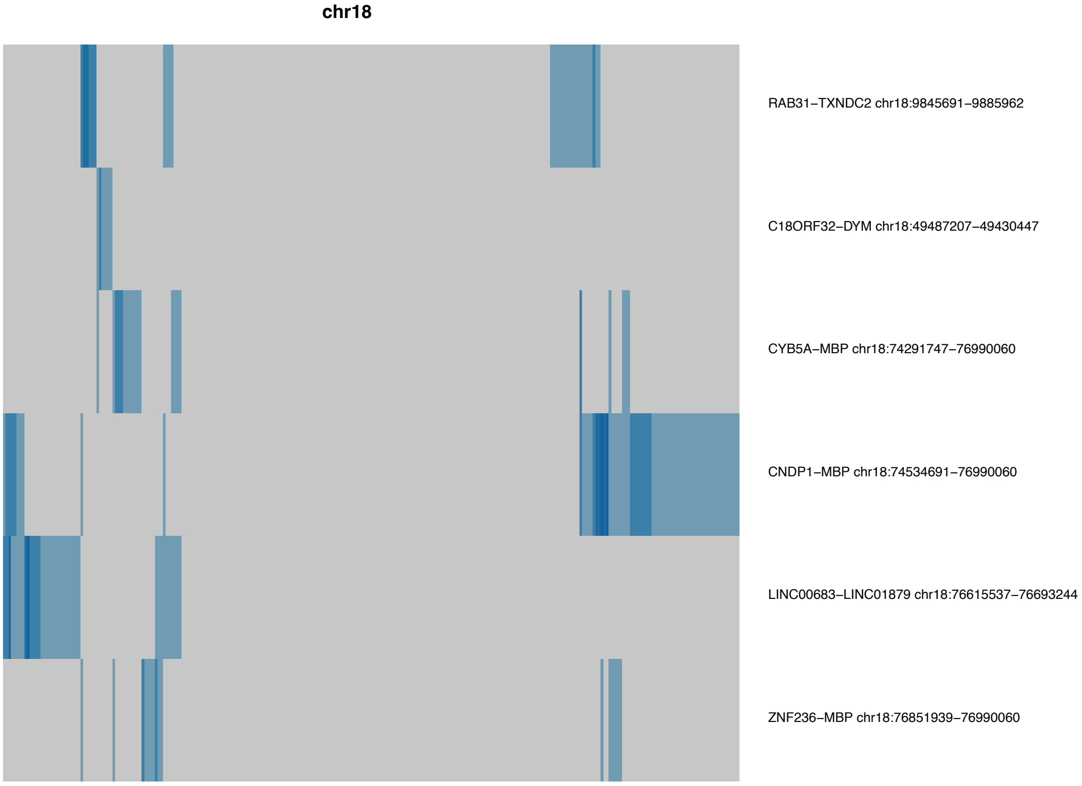


**Supplemental Figure 2. Fusion transcripts detected in short-read RNA sequencing data (all chromosomes).** Heatmaps displaying fusion transcripts detected per individual on each chromosome. Rows represent different fusion transcripts, and columns represent individuals. Depth of colour is proportional to the number of RNAseq reads supporting the fusion.

(continued next page)

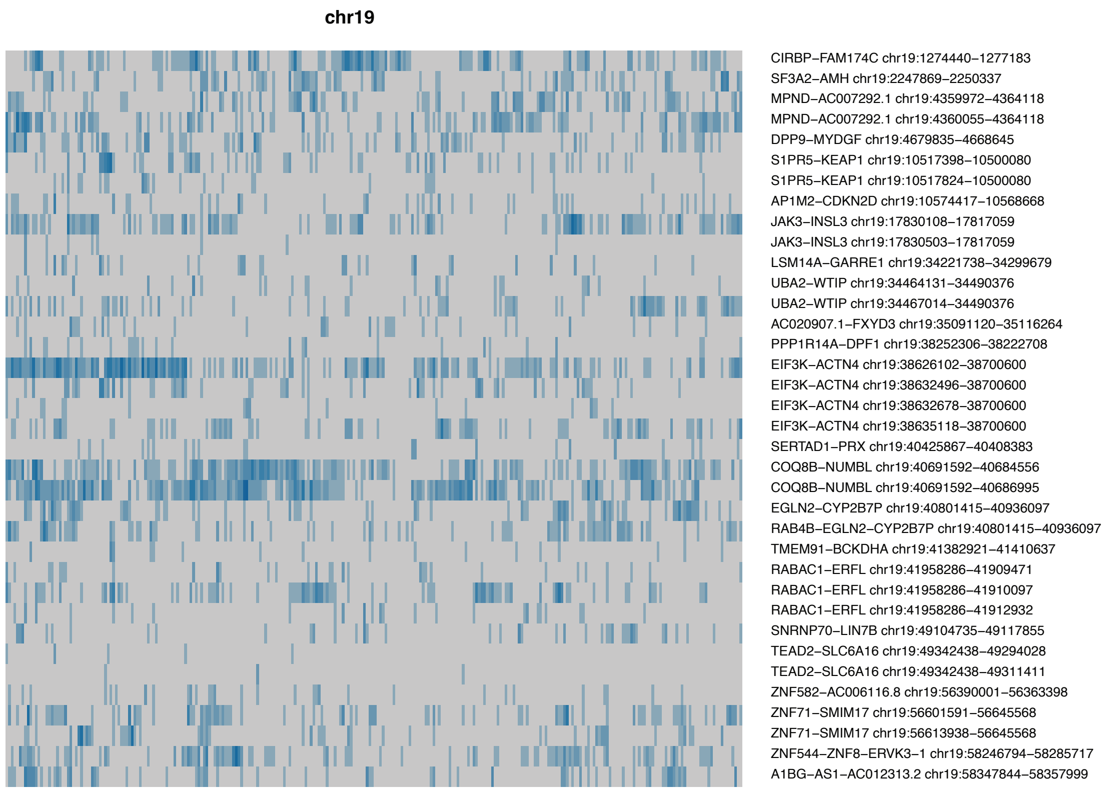


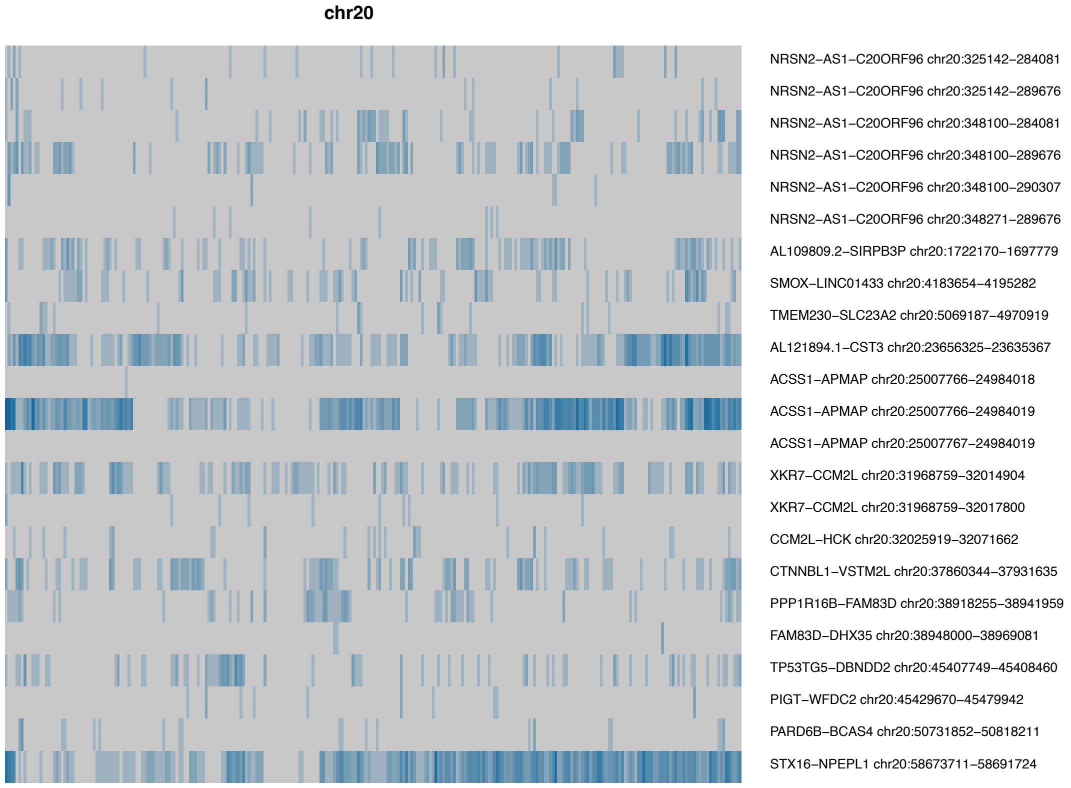


**Supplemental Figure 2. Fusion transcripts detected in short-read RNA sequencing data (all chromosomes).** Heatmaps displaying fusion transcripts detected per individual on each chromosome. Rows represent different fusion transcripts, and columns represent individuals. Depth of colour is proportional to the number of RNAseq reads supporting the fusion.

(continued next page)

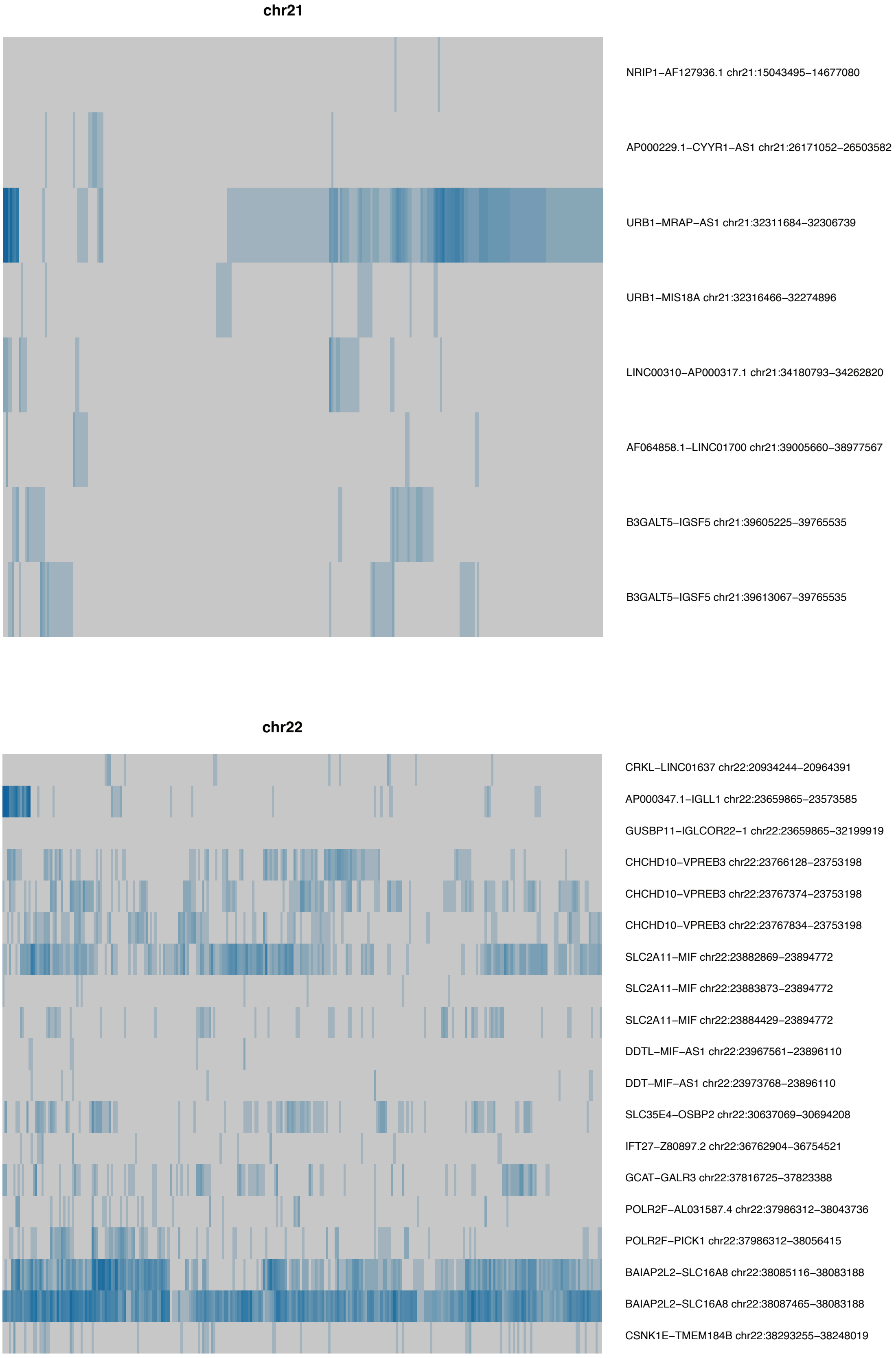


**Supplemental Figure 2. Fusion transcripts detected in short-read RNA sequencing data (all chromosomes).** Heatmaps displaying fusion transcripts detected per individual on each chromosome. Rows represent different fusion transcripts, and columns represent individuals. Depth of colour is proportional to the number of RNAseq reads supporting the fusion.

(continued next page)

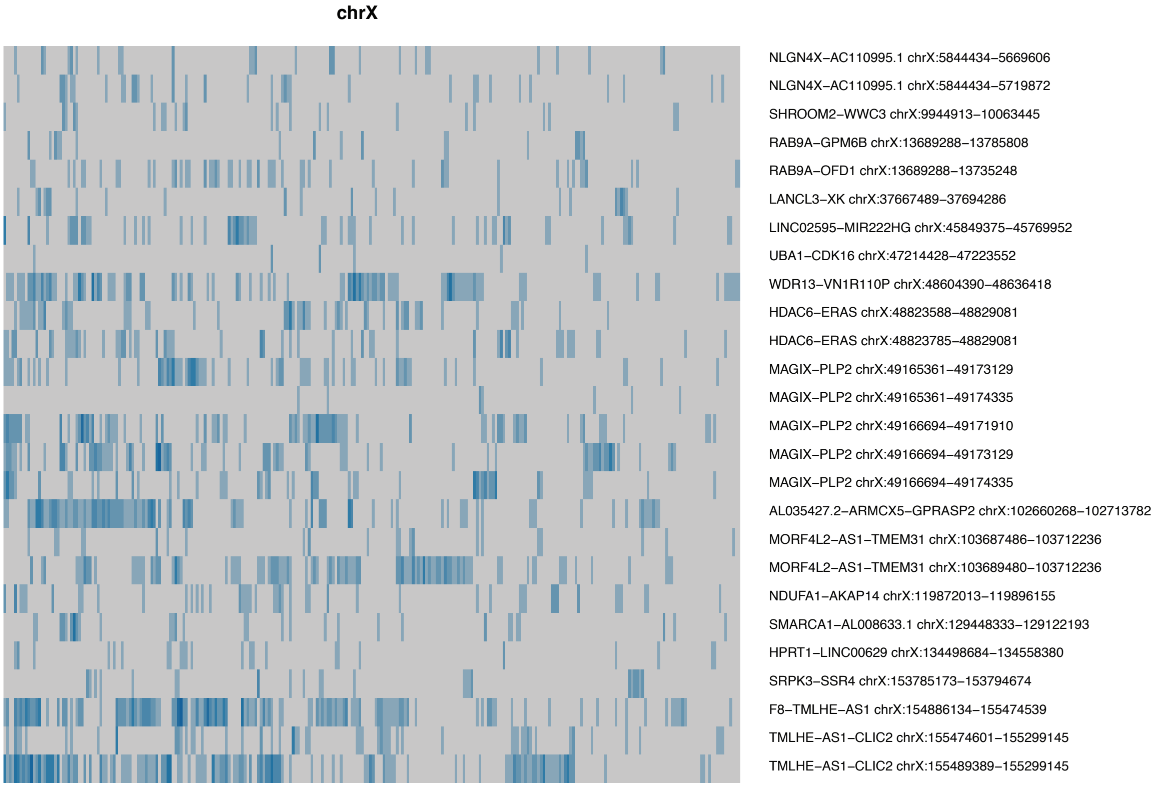


**Supplemental Figure 2. Fusion transcripts detected in short-read RNA sequencing data (all chromosomes).** Heatmaps displaying fusion transcripts detected per individual on each chromosome. Rows represent different fusion transcripts, and columns represent individuals. Depth of colour is proportional to the number of RNAseq reads supporting the fusion.

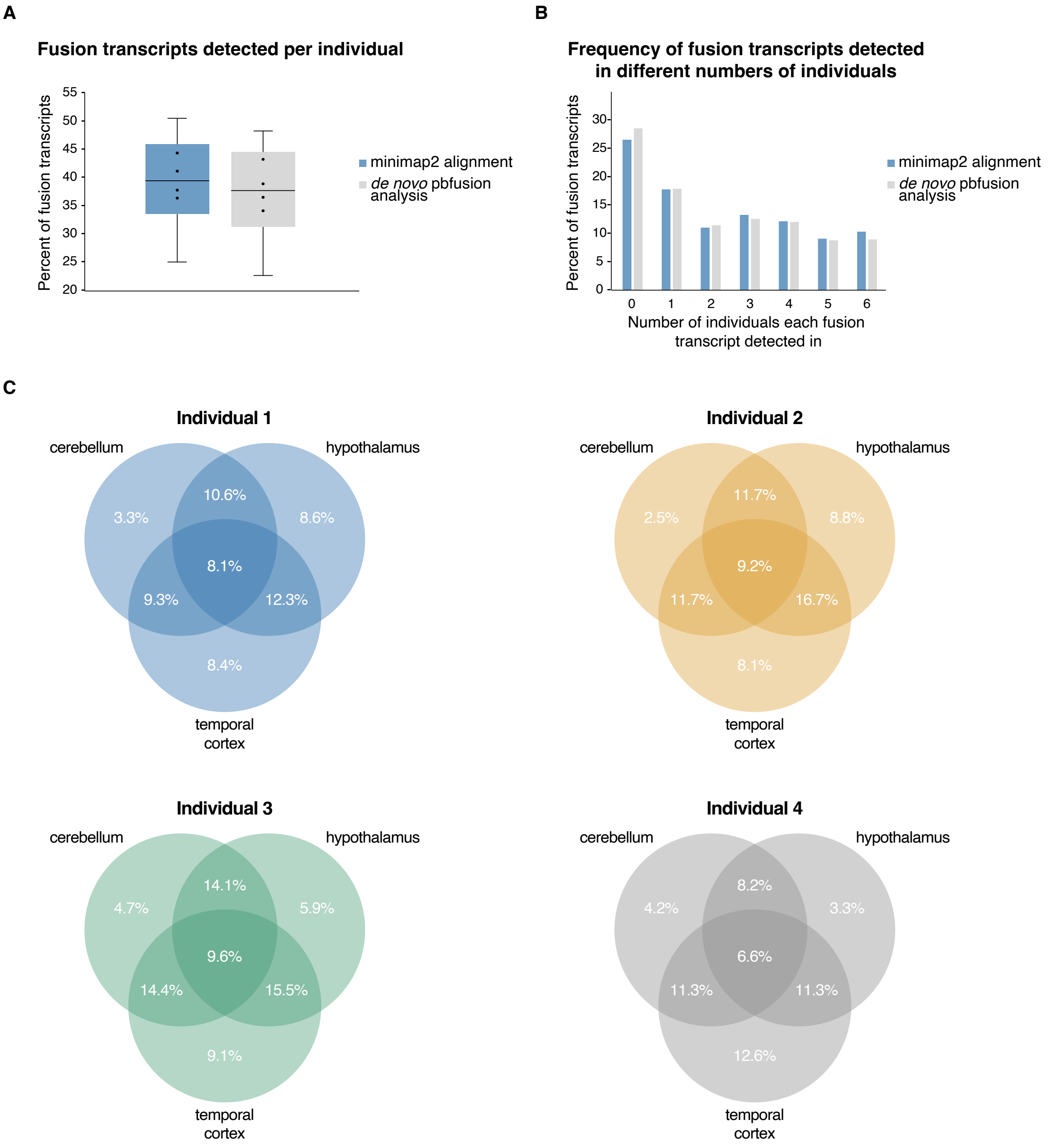


**Supplemental Figure 3. Fusion transcript validation in long-read RNA sequencing. A**: Percentage of the originally identified set of 717 fusion transcripts that were detected per individual in long-read sequencing data, using either minimap2 mapping against the fusion junction sequences, or *de novo* pbfusion fusion transcript detection. **B**: Percentage of the originally identified set of 717 fusion transcripts that were detected in different numbers of individuals in long-read sequencing data, for minimap2 mapping against the fusion junctions, and for *de novo* pbfusion fusion transcript detection; a total of 6 individuals were profiled. **C**: Overlap of fusion transcripts detected in different brain regions (cerebellum, hypothalamus, temporal cortex) from the same individual in long-read RNA-seq; n=4 individuals.

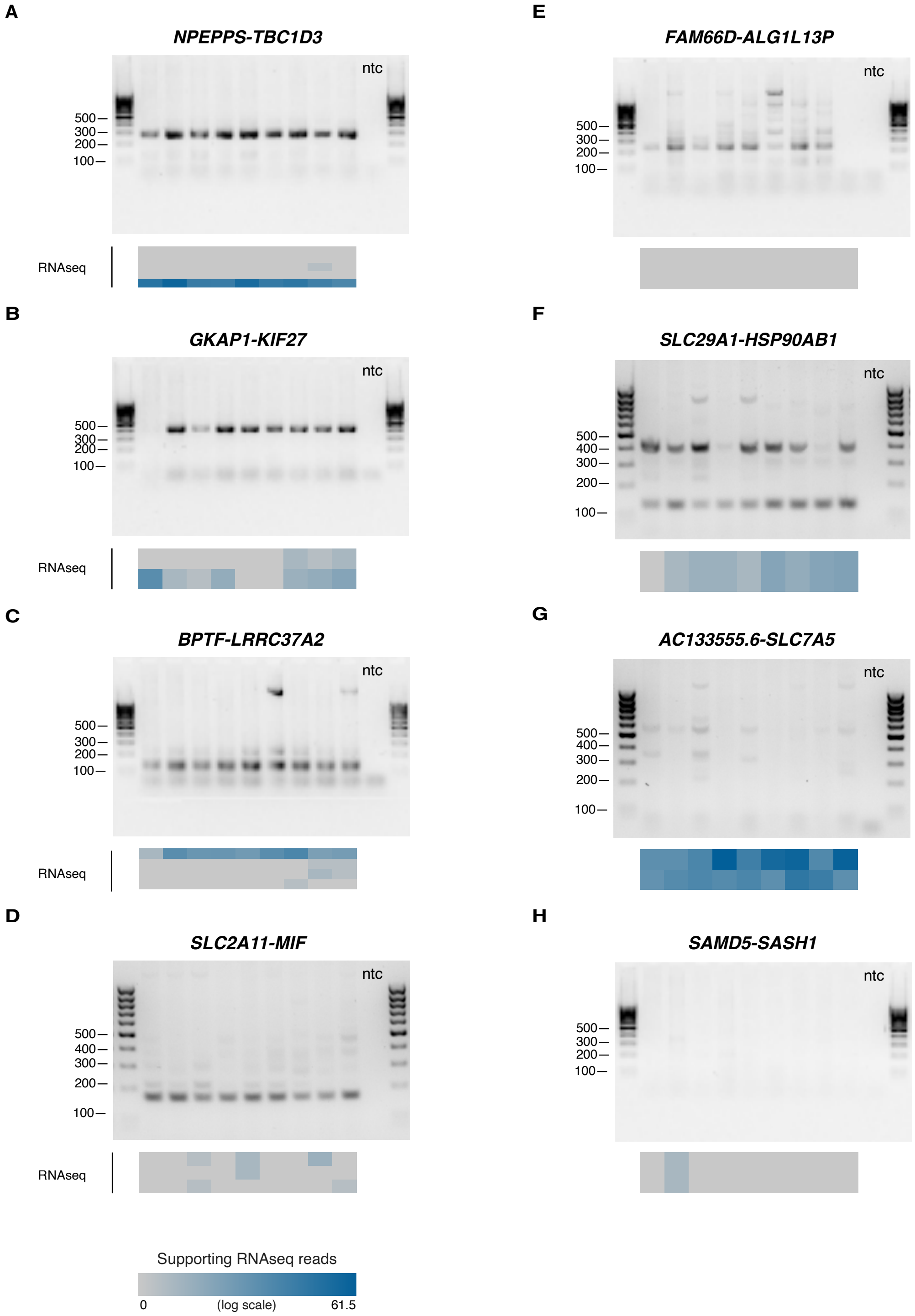


**Supplemental Figure 4. Fusion transcript expression validation in independent samples. A-H**: Gel electrophoresis analysis of RT-PCR amplified fusion transcripts from 9 individuals. Heatmaps below each gel image show the expression level of each fusion transcript based on RNA-seq analysis of the same individual. Multiple boxes in the heatmap indicate different transcript variants of the same parental genes giving rise to the fusion transcript. **A-F**: Fusion transcripts showing amplification in RT-PCR. **G-H**: Fusion transcripts that failed to amplify successfully. Ntc: no template control.

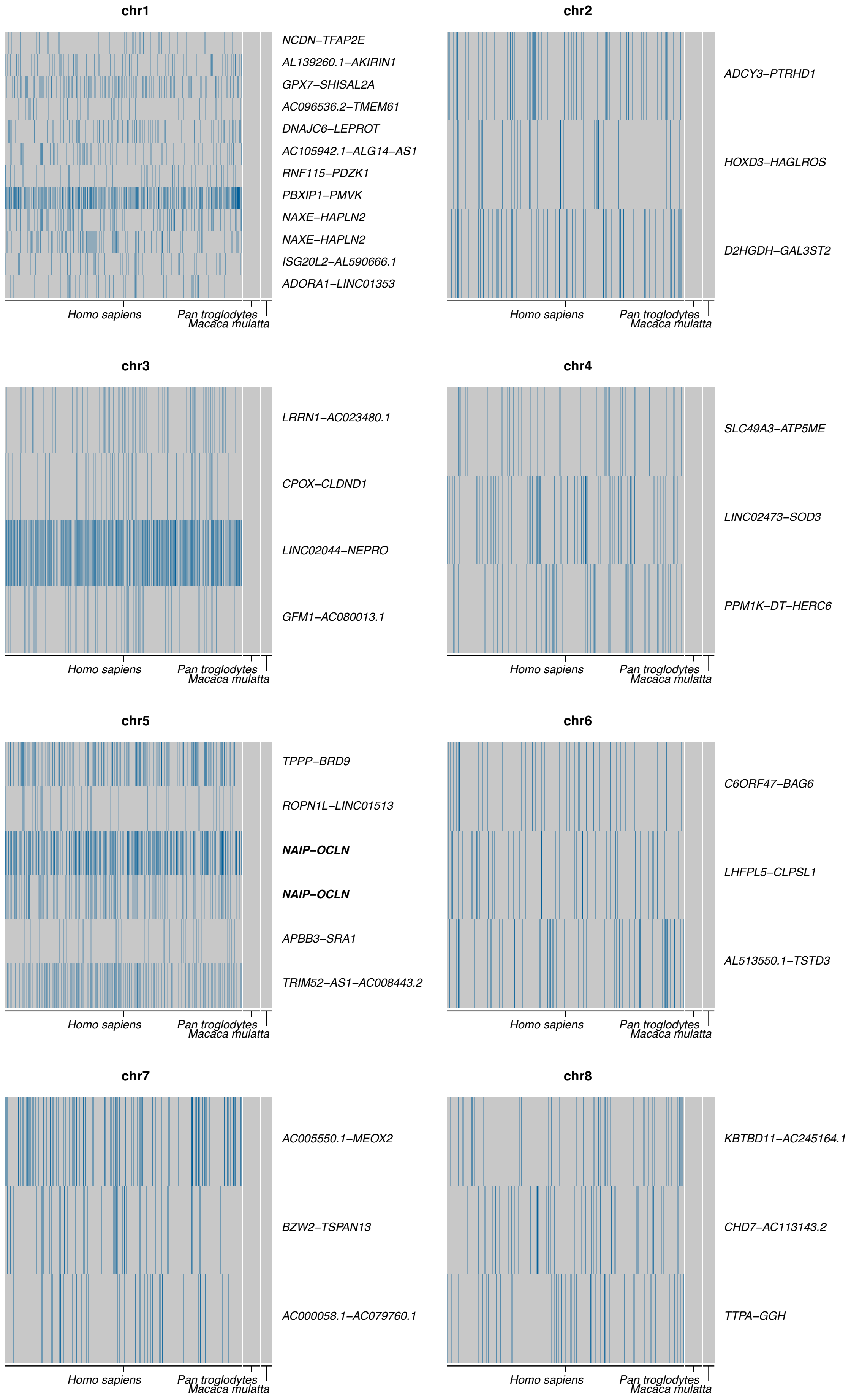


**Supplemental Figure 5. Fusion transcript detection in other primate species.** Heatmaps displaying fusion transcripts detected per individual on each chromosome, for human (*Homo sapiens*), chimpanzee (*Pan troglodytes*) and rhesus macaque (*Macaca mulatta*). Rows represent different fusion transcripts, and columns represent individuals. Depth of colour is proportional to the number of RNA-seq reads supporting the fusion. Only those fusion transcripts frequently detected in human and not detected in either of the other species ($p<0.01$) are shown.

(continued next page)

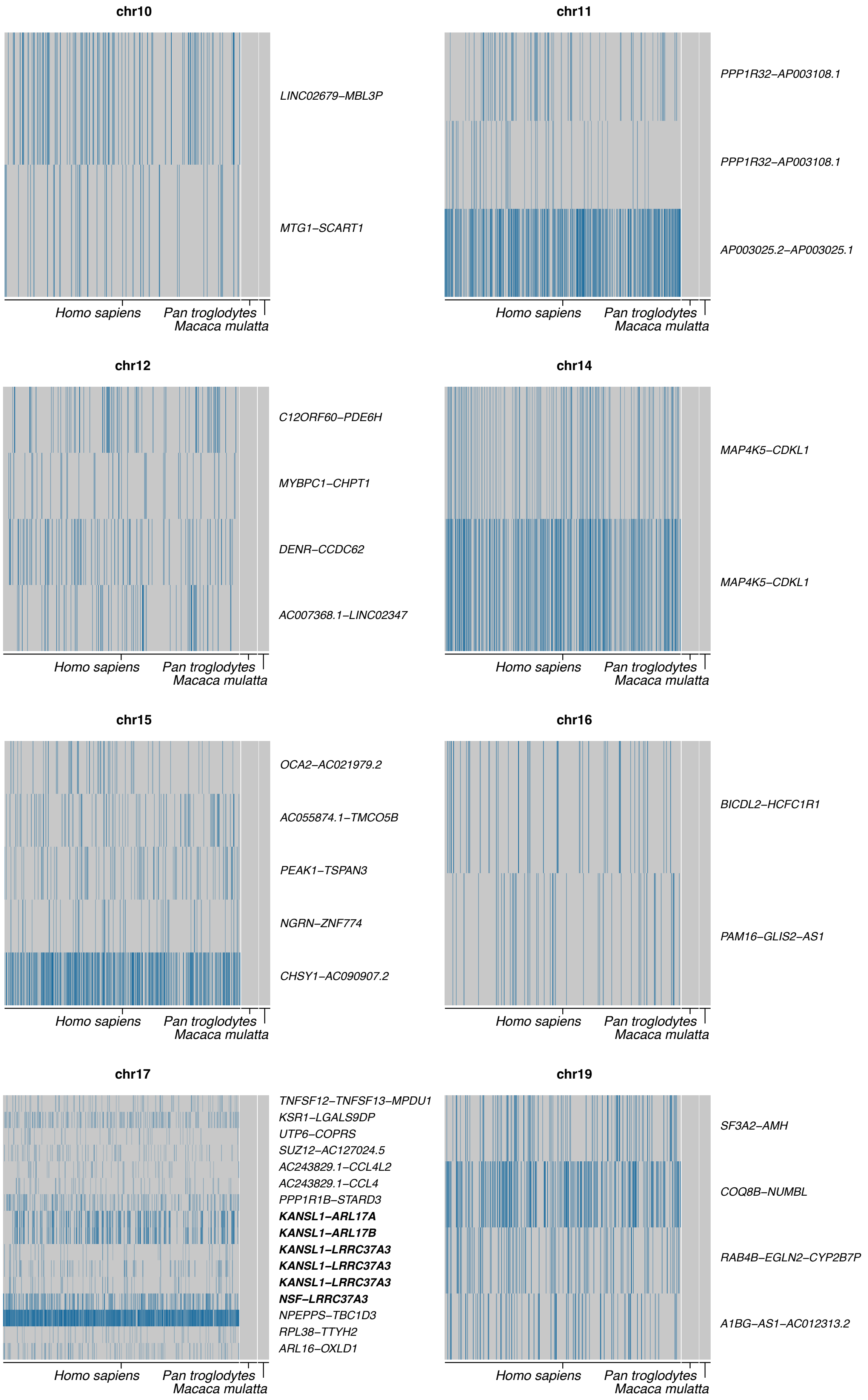


**Supplemental Figure 5. Fusion transcript detection in other primate species.** Heatmaps displaying fusion transcripts detected per individual on each chromosome, for human (*Homo sapiens*), chimpanzee (*Pan troglodytes*) and rhesus macaque (*Macaca mulatta*). Rows represent different fusion transcripts, and columns represent individuals. Depth of colour is proportional to the number of RNA-seq reads supporting the fusion. Only those fusion transcripts frequently detected in human and not detected in either of the other species ($p<0.01$) are shown.

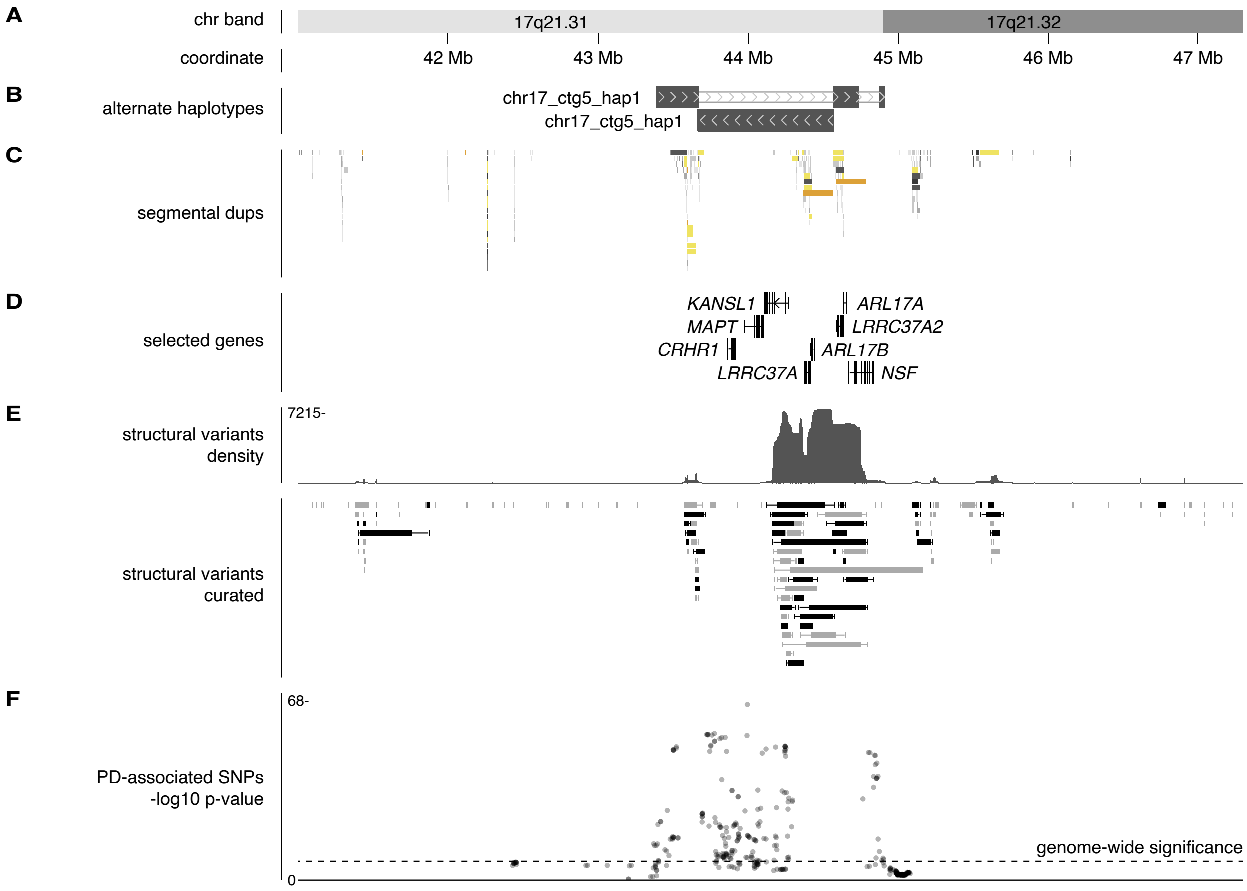


**Supplemental Figure 6. 17q21.31 is associated with Parkinson's Disease (PD) risk. A:** Chromosome band and hg38 coordinate. **B**: Location of alternate haplotypes. **C**: Segmental duplication locations. **D**: Selected genes (GENCODE v47lift37). **E**: Structural variants (deletions and duplications) found in healthy individuals from the Database of Genomic Variants (DGV). Both the density and a selection of curated variants found in multiple studies in the DGV are shown. **F**: SNPs significantly associated with PD susceptibility in previous studies; dotted line indicates genome-wide significance level.

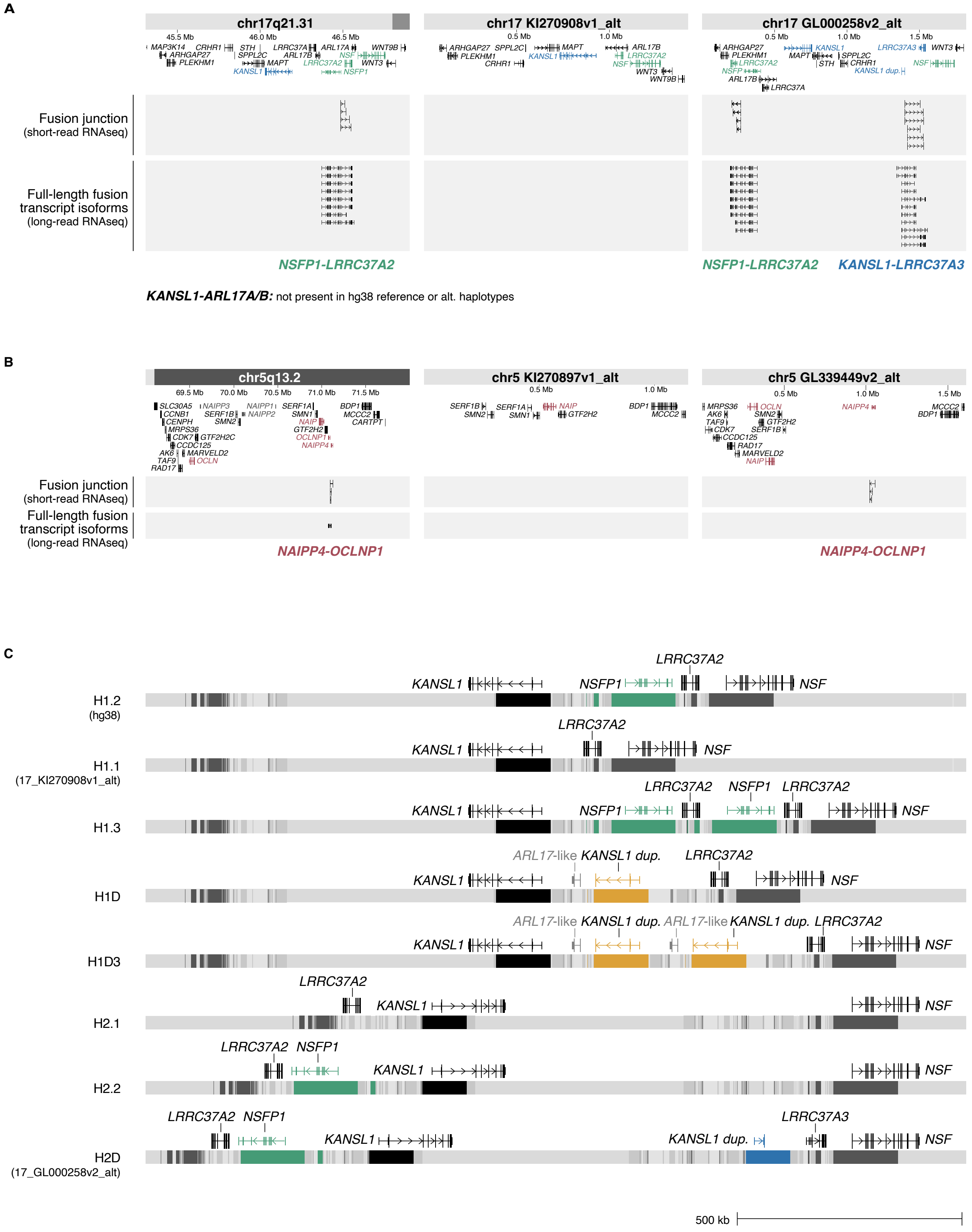


**Supplemental Figure 7. Full transcript and haplotype structures of variable fusion transcripts. A-B**: Fusion transcript isoform structures discovered from long-read Pacbio and Oxford Nanopore Technologies sequencing, mapped to hg38 and alternate haplotypes for 17q21.31 (**A**) and 5q13.2 (**B**). Full-length parental genes and the partially duplicated genes that originated from them are highlighted in colour. Fusion junctions identified from FusionCatcher short-read RNA-seq analysis and full fusion transcript isoform structure identified from Pacbio sequencing and isoform discovery are shown. *KANSL1-ARL17A/B* was not mapped to hg38 or any alternate haplotypes as the structural haplotype it originates from, H1D, is not represented in the human reference genome. **C:** Structure of 17q21.31 genome region indicating proposed location of genes generating fusion transcripts. All eight structural haplotypes are shown (based on Steinberg et al., 2012 and Boettger et al., 2012). Coloured boxes represent segmental duplications containing partially duplicated genes that give rise to fusion transcripts. The locations of full-length and partially duplicated genes giving rise to fusion transcripts are displayed, and the duplicated genes causing the fusion event are shown in colour. H1.2, H1.1, H1.3, H1D, H1D3, H2.1, H2.2 and H2D refer to structural haplotype nomenclature of Steinberg et al. (2012). Haplotype H1.2 is represented as the hg38 reference genome, while H1.1 and H2D are represented as alternate haplotypes 17_KI270908v1_alt and chr17_GL000258v2_alt, respectively. The other structural haplotypes are not represented in hg38 or alternate haplotypes.

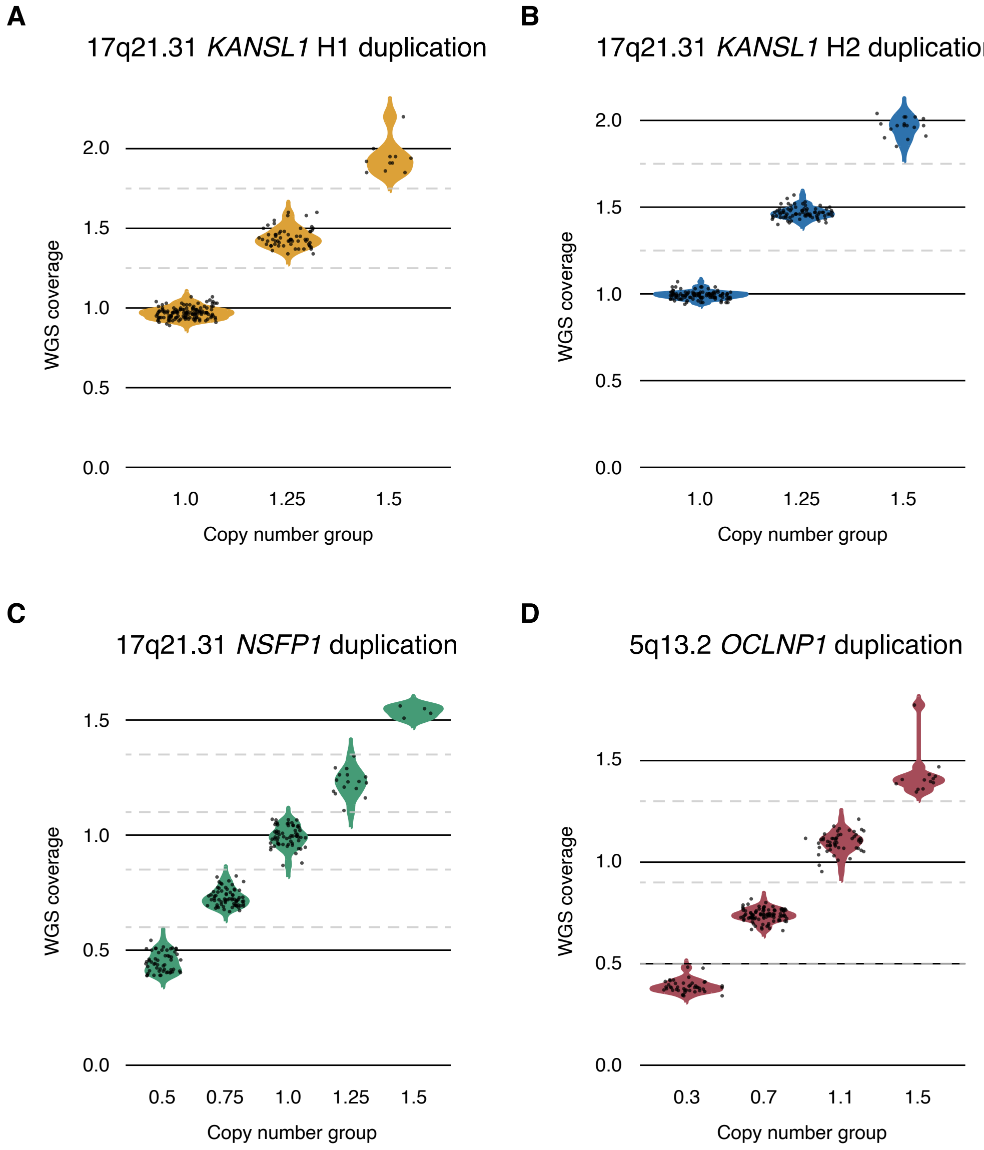


**Supplemental Figure 8. Individuals showing varying copy number of segmentally duplicated regions at 17q21.31 and 5q13.2. A-D**: Whole genome sequencing coverage for duplication regions at 17q.21.31 and 5q13.2, expressed as a ratio relative to nearby single-copy, non-duplicated regions; n=191 individuals. Each duplication contains a partially duplicated gene giving rise to variable fusion transcripts. Individuals were separated into copy number groups (shown on the *x*-axis) for downstream analysis of fusion transcript expression in each group.

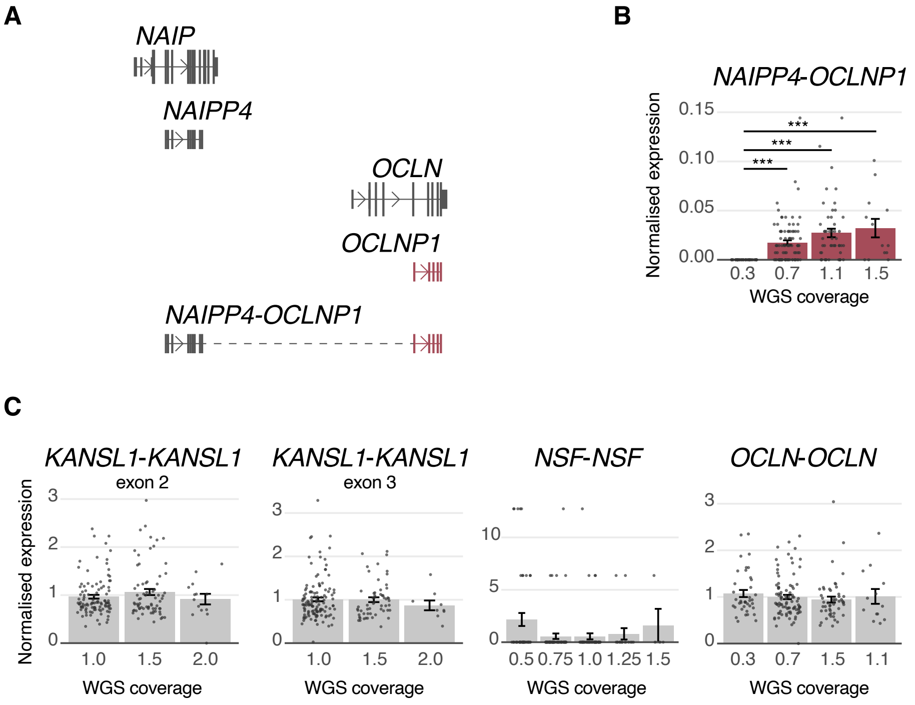


**Supplemental Figure 9. Variation in *NAIPP4-OCLNP1* expression arises from different genomic structural haplotypes. A**: Structure of *NAIPP4-OCLNP1* fusion transcript, based on Illumina, Pacbio and Oxford Nanopore Technologies sequencing data. Not all fusion transcript splice variants are depicted here; additional transcript variants are displayed in **Supplemental Figure 6**. **B**: Expression level of fusion transcript based on genomic copy number of relevant segmental duplication. Segmental duplication copy number is indicated by the whole genome sequencing coverage level. **C**: Expression level of full-length parental genes giving rise to the partially duplicated genes that generate variable fusion transcripts. Fusion transcript and parental control transcript expression are normalised to the mean parental control transcript expression level; n=191 individuals.

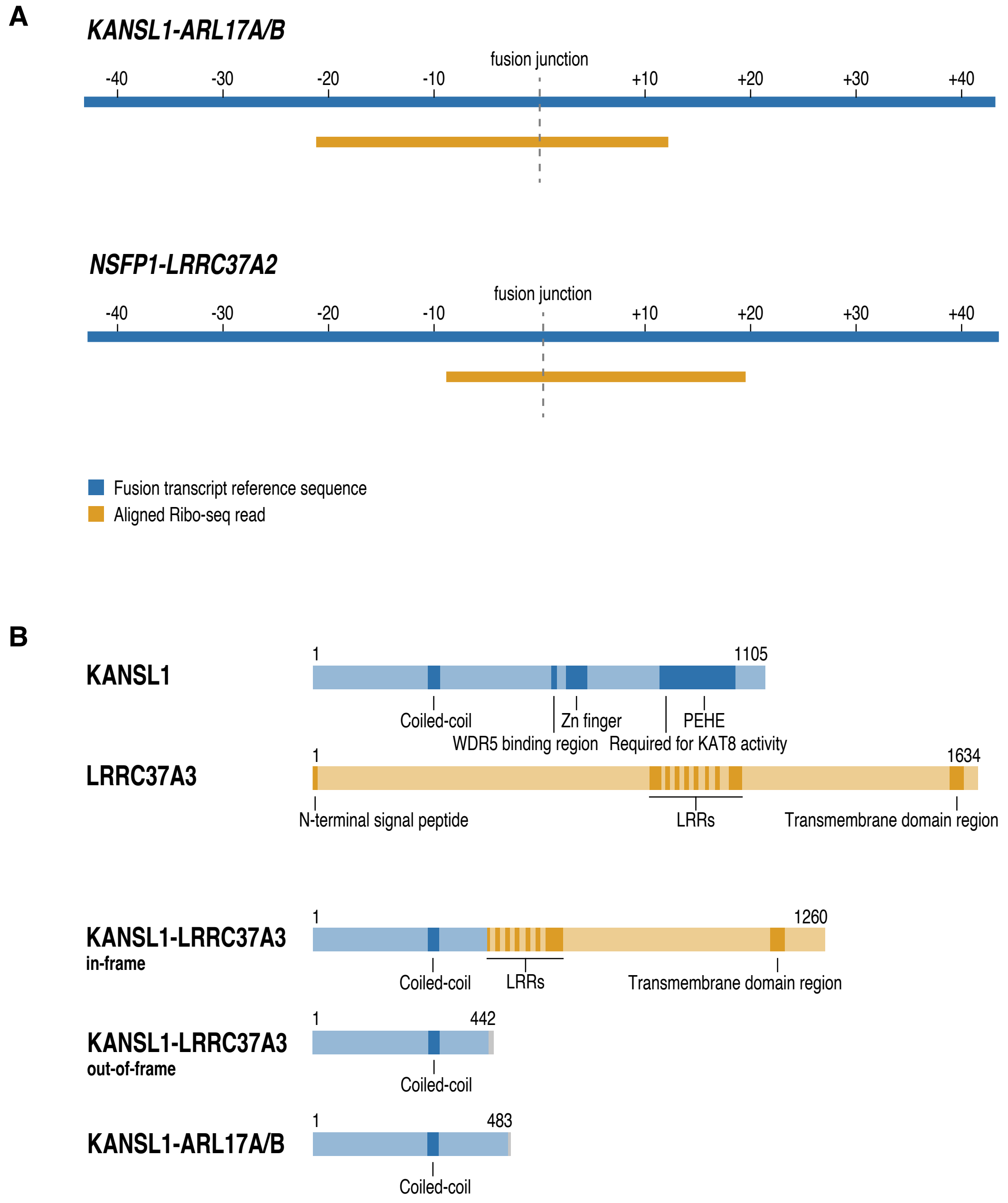


**Supplemental Figure 10. Fusion proteins encoded by variable fusion transcripts. A**: Supporting Ribo-seq reads for *KANSL1-ARL17A/B* and *NSFP1-LRRC37A2*. **B**: Predicted protein structures for KANSL1-LRRC37A3 in-frame and out-of-frame variants, KANSL1-ARL17A/B and NSFP1-LRRC37A2. Functional domains are labelled.
